# Polyadenylation of Histone H3.1 mRNA Promotes Cell Transformation by Displacing H3.3 from Gene Regulatory Elements

**DOI:** 10.1101/806828

**Authors:** Danqi Chen, Qiao Yi Chen, Zhenjia Wang, Yusha Zhu, Thomas Kluz, Wuwei Tan, Jinquan Li, Feng Wu, Lei Fang, Xiaoru Zhang, Rongquan He, Steven Shen, Hong Sun, Chongzhi Zang, Chunyuan Jin, Max Costa

**Affiliations:** Department of Environmental Medicine, New York University School of Medicine, New York, NY, USA; Department of Biochemistry and Molecular Pharmacology, New York University School of Medicine, New York, NY, USA; Perlmutter Cancer Center, NYU Langone Health, New York, NY, USA; Center for Public Health Genomics, University of Virginia School of Medicine, Charlottesville, VA, USA; Department of Statistics, University of Virginia, Charlottesville, VA, USA; Hubei Province Key Laboratory of Occupational Hazard Identification and Control, Medical college, Wuhan University of Science and Technology; Medical School of Nanjing University, Nanjing, China; Department of Medical Oncology, First Affiliated Hospital of Guangxi Medical University, Nanning, China; Institute of Health Informatics, University of Minnesota, Minneapolis, MN, USA; Department of Public Health Sciences, University of Virginia, Charlottesville, VA, USA; Department of Electrical and Computer Engineering, Texas A&M University, College Station, TX, USA

**Keywords:** Canonical histones, Histone variants, H3.1, H3.3, Polyadenylation, Arsenic

## Abstract

Replication-dependent canonical histone messenger RNAs (mRNAs) do not terminate with a poly(A) tail at the 3’ end. We previously demonstrated that exposure to arsenic, an environmental carcinogen, induces polyadenylation of canonical histone H3.1 mRNA. The addition of a poly(A) tail to the H3.1 mRNA caused transformation of human cells *in vitro*, but the underlying mechanisms are unknown. Here we report that polyadenylation of H3.1 mRNA increases H3.1 protein level, resulting in depletion of histone variant H3.3 at active promoters, enhancers, and insulator regions through its displacement. Cells underwent transcriptional deregulation, G2/M cell cycle arrest, chromosome aneuploidy and aberrations. Furthermore, knocking down the expression of H3.3 induced cell transformation, whereas ectopic expression of H3.3 attenuated arsenic-induced cell transformation, suggesting that H3.3 displacement might be central to tumorigenic effects of polyadenylated H3.1 mRNA. Our study provides novel insights into the importance of proper histone stoichiometry in maintaining genome integrity.

**Highlights:**

- Polyadenylation of canonical histone H3.1 mRNA promotes tumor formation in nude mice
- Histone variant H3.3 is displaced from critical gene regulatory elements by overexpression of polyadenylated H3.1 mRNA
- Increased polyadenylated H3.1 mRNA causes abnormal transcription, cell cycle arrest, and chromosomal instability
- Arsenic induces polyadenylation of H3.1 mRNA *in vivo*

## Introduction

Histone genes are highly conserved through evolution. While replication-dependent histones, referred to as canonical, constitute the majority of histone genes, various replication-independent histone genes which encode variants have evolved. Each variant retains distinct locus-specific properties and is unique in terms of gene and protein sequences [1]. Canonical histone genes (e.g., histone H3.1) lack introns and are the only genes found in multicellular organisms whose messenger RNAs (mRNAs) that do not terminate with a poly (A) tail at their 3’ end [2]. Instead, canonical histone mRNAs terminate with a stem-loop structure, which is comprised of a highly conserved 26-nucleotide sequence that binds Stem-loop binding protein (SLBP), an important component for canonical histone pre-mRNA processing and translation [3].

Canonical histone mRNAs are rapidly degraded at the end of S phase probably due to the lack of a poly (A) tail at the 3’ end, rendering them with decreased mRNA stability [3, 4]. Thus, lack of poly(A) tail is vital for maintaining the steady state levels of canonical histone mRNAs during S phase in normal cells. In the event of pre-mRNA processing disruption, for instance through loss of cellular SLBP, canonical histone mRNAs are greatly stabilized by gaining a poly (A) tail, resulting in excess canonical histones [5, 6]. This could disrupt appropriate histone stoichiometry and induce genome instability. Indeed, in *Drosophila*, deletion and mutation of SLBP have been found to induce polyadenylation of canonical histone mRNAs and cause zygotic lethality partially by a failure of chromosome condensation that blocks normal mitosis process [5–8]. However, whether and how polyadenylated canonical histone mRNAs affect genomic stability and other cellular processes have not been studied.

We previous demonstrated that 3’ end of canonical histone H3.1 mRNA can be polyadenylated by exposure to an environmental carcinogen arsenic [9]. Arsenic causes human lung, bladder, and skin cancers [10] and is linked to cancers of the kidney, liver, and prostate [11–13]. In an attempt to examine tumorigenic effect of polyadenylated H3.1 mRNA, we carried out the soft agar colony formation assays. Overexpression of polyadenylated H3.1 mRNA enhanced anchorage-independent growth of human bronchial epithelial BEAS-2B cells, which indicated that polyadenylation of H3.1 mRNA possesses capacity to induce cell transformation [14].

Polyadenylated canonical histone H3.1 mRNA might induce genomic instability and eventually cell transformation by disrupting the balance between canonical and variant histones. Deposition of canonical histone H3.1 is coupled with DNA replication and its expression is largely limited to S phase of the cell cycle, while histone variant H3.3 is expressed throughout the cell cycle in a DNA replication-independent manner [15–19]. Polyadenylated canonical mRNAs are not cell cycle regulated, for instance, arsenic-induced addition of a polyadenylated tail allowed expression of H3.1 outside of S phase [9]. Therefore, it is conceivable that accumulated or mistimed expression of H3.1 during the cell cycle may affect H3.3 assembly that could have significant impacts on chromatin landscape and genome integrity since H3.3 has unique properties and functions distinct from canonical H3 [1, 20].

This study investigates the mechanisms that underlie tumorigenic effects of polyadenylated H3.1 mRNA. We find that polyadenylated H3.1 mRNA displaces the variant H3.3 from important gene regulatory elements such as active promoters, enhancers, and insulator regions. Reducing H3.3 expression, which mimics H3.3 depletion by H3.1 polyadenylation, induces cell transformation, while overexpression of H3.3 rescues cell transformation induced by arsenic. H3.3 displacement thus appears to be central to tumorigenic effects of polyadenylated H3.1 mRNA. Our findings also add important insights not only into genomic instability induced by imbalance in histone stoichiometry but also into the oncogenic role for H3.3.

## Results

### Polyadenylation of canonical histone H3.1 mRNA enhances tumor formation in nude mice

Our previous study has demonstrated that arsenic induces polyadenylation of canonical histone H3.1 mRNA, through depletion of SLBP in tissue culture models [9]. To study the impact of arsenic-induced polyadenylation of canonical histone H3.1 mRNA on carcinogenesis, we analyzed the effect of H3.1 mRNA polyadenylation on cell proliferation, anchorage-independent growth, and tumor formation *in vivo*. We first constructed the H3.1poly(A) plasmid, containing a poly(A) signal immediately downstream of a H3.1 cDNA fragment to express polyadenylated histone H3.1 mRNA (Figure 1A). Since antibodies against H3 are not able to distinguish different types of histone H3, human bronchial epithelial cells (BEAS-2B) were stably transfected with FLAG-tagged H3.1poly(A) plasmid (FH3.1poly(A)) (Figure 1A). Overexpression of both total and polyadenylated H3.1 mRNA was confirmed using RT-qPCR (Figure 1B). The amount of total and polyadenylated mRNA was measured using cDNAs synthesized with random or oligo(dT) primers, respectively (Figure 1B). Total H3 protein levels were increased by 2.67-fold as compared with the control (compare lanes 1 and 2 in Figure 1C; 1.90+2.03/1.47=2.67 vs. 1/1=1), which is comparable to the two-fold increase induced by arsenic exposure [9]. Thus, results obtained with the FH3.1poly(A) plasmid are representative of arsenic-induced H3.1 polyadenylation.

**Figure 1.**
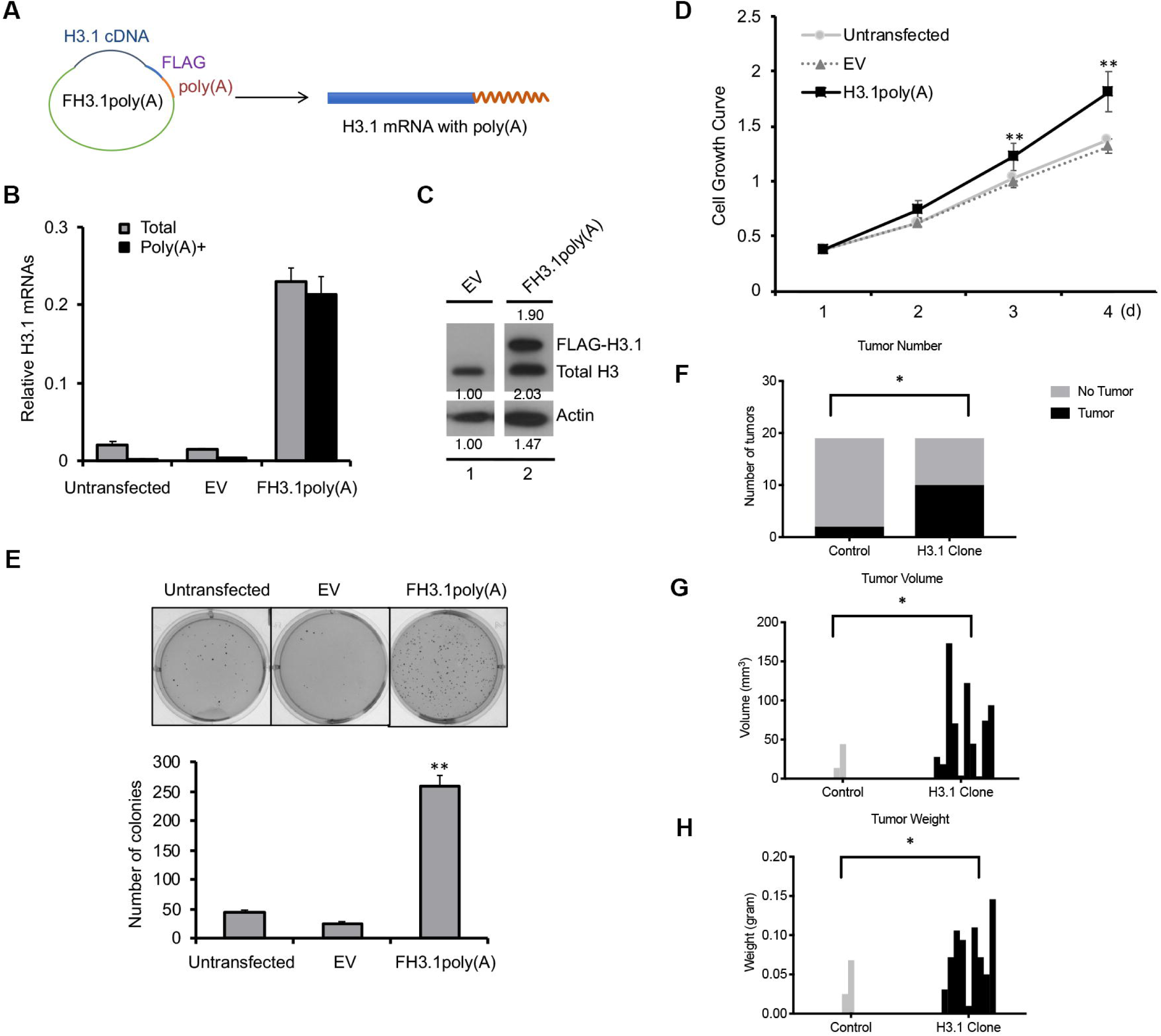
Polyadenylation of canonical histone H3.1 mRNA enhances tumor formation in nude mice. **(A)** A H3.1 cDNA fragment was inserted into the multiple clone sites (MCSs) of pcDNA-FLAG vector, which contains a poly(A) signal in the downstream of MCSs and generates polyadenylated mRNAs. **(B)** pcDNA-FLAG-Empty (EV) or FH3.1poly(A) vector were stably transfected into BEAS-2B cells. The amount of total and polyadenylated H3.1 mRNA was measured by RT-qPCR using cDNAs synthesized with random or oligo(dT) primers, respectively. **(C)** Western blot shows exogenous H3.1 protein generated by transfection of FH3.1poly(A). **(D)** Cell growth rate was measured by MTT. The data shown are the mean ±S.D. (n=3). ** *p*<0.01 **(E)** Soft agar assays. The data shown are the mean ±S.D. (n=3). ** *p*<0.01. **(F-H)** A total of 5 million transformed or control cell clones were injected into nude mice. After 5 months post-injection, the mice were sacrificed. Tumor number (F), volume (G), and weight (H) were calculated. Tumor diameters were measured with calipers, and the tumor volume was calculated. Results represent the mean ±S.D. (n=18). * *p*<0.05.

The rate of cell proliferation was increased in BEAS-2B cells transfected with FH3.1poly(A) plasmid compared to cells transfected with empty vector (Figure 1D). Consistent with previous results [14], BEAS-2B/FH3.1poly(A) cells displayed enhanced anchorage-independent growth compared with BEAS-2B cells transfected with or without empty vector (Figure 1E). To evaluate the tumorigenic effect of polyadenylated H3.1 mRNA *in vivo*, we next collected soft agar clones formed by control and FH3.1poly(A)-expressing BEAS-2B cells, respectively, and injected them into nude mice to assess tumor formation. Juxtaposing the two groups, mice injected with FH3.1poly(A) cells exhibited 50% tumor formation compared to 11% in the control group (Figure 1F). Moreover, xenograft of FH3.1poly(A) cells in mice displayed significantly higher volume and weight than the control (Figure 1G and H). Taken together, these data suggest that polyadenylation of H3.1 mRNA is able to promote tumor cell growth in nude mice.

### Insertion of a stem-loop sequence before the poly(A) signal attenuates cell transformation induced by polyadenylation of H3.1 mRNA

To examine whether acquisition of a poly(A) tail is the determining factor for cell transformation induced by transfection of FH3.1poly(A) plasmid, we constructed a plasmid H3.1Loop containing a stem-loop sequence residing in front of the poly(A) signal (Figure 2A). Following binding of SLBP and other 3’ pre-mRNA processing factors to the stem-loop sequence, a majority of transcriptions are terminated immediately downstream of the sequence, before the RNA polymerase can reach the poly(A) signal. Thus, the presence of the stem-loop sequence is expected to generate mRNAs with less poly(A) tails (Figure 2A). To rule out potential effects resulting from FLAG tag (Figure 1), we prepared H3.1poly(A) and H3.1Loop plasmids without any tag (Figure 2A). RT-qPCR results demonstrated that transfection with the plasmid containing H3.1Loop sequence significantly reduced the amount of polyadenylated H3.1 mRNA compared with cells transfected with H3.1poly(A) vector (Figure 2B). Moreover, western blot analyses revealed lower levels of H3 protein in cells transfected with H3.1Loop compared with H3.1poly(A) (Figure 2C). The reduction of H3 protein in H3.1Loop-transfected cells was likely due to the instability and degradation of un-polyadenylated histone mRNAs. The difference seemed not to have resulted from the variation of inserted DNA copy numbers, since PCR results showed that the level of H3.1Loop DNA in H3.1Loop-transfected cells was comparable to the level of H3.1polyA DNA in H3.1poly(A)-transfected cells (Figure 2D). Taken together, any differential results obtained from using H3.1poly(A) and H3.1Loop will likely be attributable to the effects of mRNA polyadenylation.

**Figure 2.**
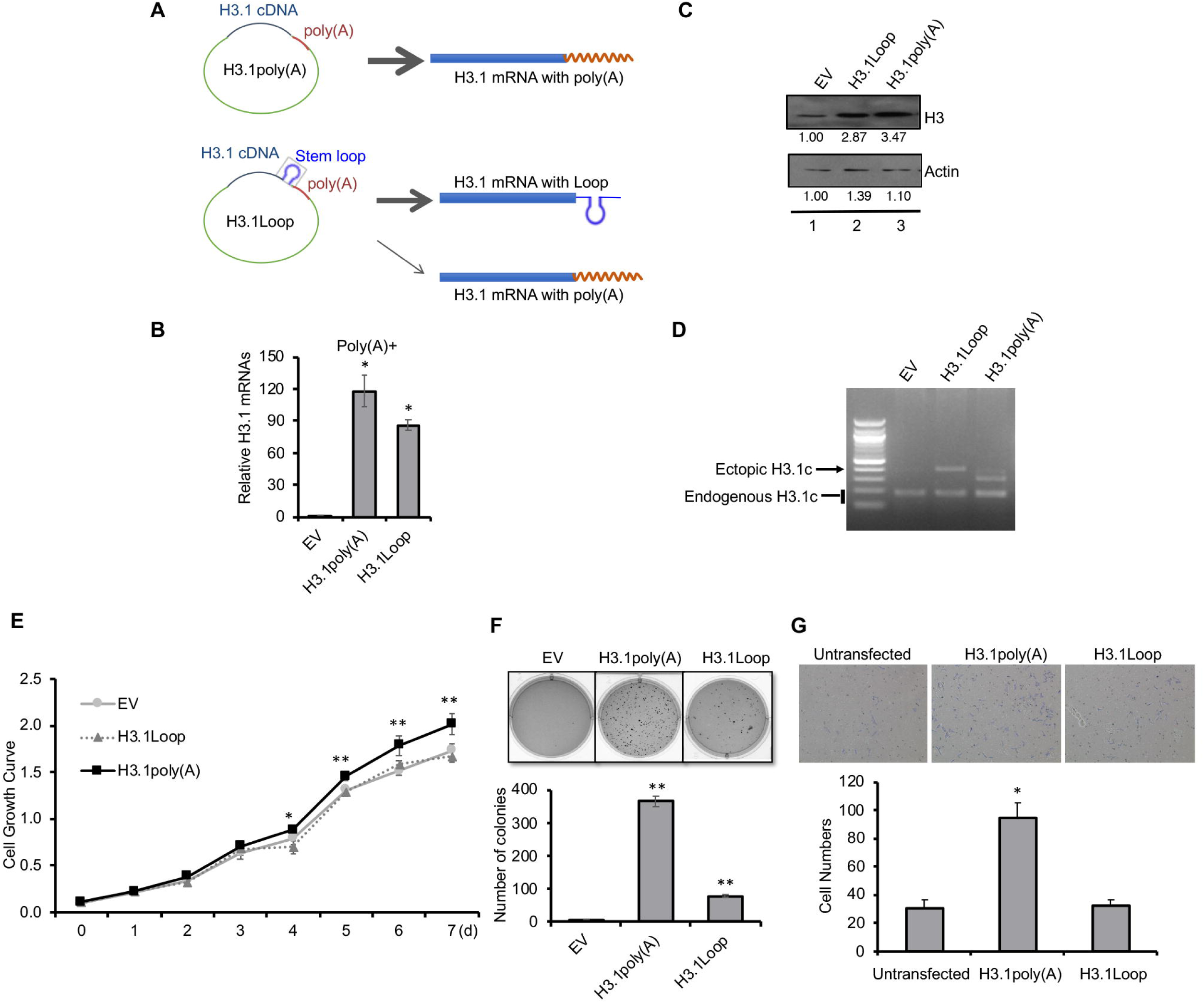
Insertion of a stem-loop sequence before the poly(A) signal attenuates cell transformation induced by polyadenylation of H3.1 mRNA. **(A)** A H3.1 cDNA fragment was inserted into the MCSs of pcDNA vector, which contains a poly(A) signal in the downstream of MCSs. H3.1Loop plasmid: a stem-loop sequence was inserted before the poly(A) signal, whose transcription terminates near the stem-loop sequence, generating H3.1 mRNAs mostly without poly(A) tail. pcDNA-Empty (EV), H3.1poly(A) or H3.1Loop vector was stably transfected into BEAS-2B cells. **(B)** RT-qPCR results using oligo (dT) primers for reverse transcription to capture polyadenylated H3.1 mRNAs. **(C)** Western blot analysis. The band intensities were quantified using ImageJ software. The controls in lane 1 were used as references (set to 1). **(D)** Total DNA was extracted and exogenous H3.1 was then amplified by PCR with primers for H3.1c and pcDNA vector, while endogenous H3.1 was amplified with primers specific for H3.1c. **(E)** The cell growth rate was measured by MTT. The data shown are the mean ±S.D. (n=3). \**p*<0.05; ** *p*<0.01. **(F)** Soft agar assays. The data shown are the mean ±S.D. (n=3). ** *p*<0.01. **(G)** Transwell invasion assays. The migrated cells were fixed, stained, and counted. Bars indicate S.D. (n=3). * *p*<0.05.

We next examined how transfection of H3.1poly(A) and H3.1Loop differentially affected cell growth, transformation, and invasion. While H3.1poly(A)-transfected cells demonstrated accelerated growth compared with controls, transfection with H3.1Loop reduced the rate of cell growth to the same level as the control (Figure 2E). Anchorage-independent cell growth was enhanced by the transfection of H3.1poly(A) as compared with control, an observation consistent with the result seen with transfection of FLAG-tagged H3.1poly(A) (Figure 1E), suggesting that FLAG tag was not the cause of the observed cell transformation. Notably, insertion of the Loop sequence upstream of the poly(A) signal greatly attenuated anchorage-independent cell growth (Figure 2F). Likewise, H3.1poly(A) cells exhibited much higher rate of cell migration than H3.1Loop and control cells in Transwell invasion assays (Figure 2G). The above results indicate the critical function of polyadenylation of mRNA in cell transformation and invasion induced by ectopic expression of canonical histone H3.1 mRNA.

### Overexpression of polyadenylated H3.1 mRNA displaces histone variant H3.3 from critical gene regulatory elements

Polyadenylation of H3.1 mRNA results in the increase in H3.1 protein level (Figure 1C and Figure 2C). Moreover, we have previously demonstrated that arsenic exposure induces polyadenylation of canonical histone H3.1 mRNA, resulting in the accumulation of H3.1 mRNA in Mid-to late S phase and M phase [9]. The expression of canonical histone H3.1 peaks during S phase, while variant H3.3 is expressed throughout the cell cycle. Thus, both the increase in H3.1 protein and the presence of H3.1 protein outside of S phase resulting from polyadenylation of H3.1 mRNA may have an impact on the assembly of variant histone H3.3, which differs from H3.1 by only five amino acids. We used chromatin immunoprecipitation followed by high-throughput sequencing (ChIP-seq) to examine the potential effects of polyadenylated H3.1 mRNA on H3.3 assembly. BEAS-2B cells stably expressing FLAG-tagged H3.3 were transfected with untagged H3.1poly(A) plasmid or empty vector. RT-qPCR results confirmed the overexpression of polyadenylated H3.1 mRNA (Figure S1A). Notably, the total cellular level of FLAG-H3.3 was not changed by ectopic expression of H3.1poly(A) as shown in Figure S1B.

H3.3-containing nucleosomes were isolated by anti-FLAG affinity-gel purification followed by ChIP-seq to examine how genome-wide H3.3 deposition was influenced by ectopic expression of H3.1poly(A). Results revealed highly enriched H3.3 in the promoter regions of 2,000 of the most highly expressed genes in control cells (blue line, Figure 3A). This is consistent with our previous results seen in another cell line [21]. The deposition of H3.3, however, was greatly reduced in the cells transfected with untagged-H3.1poly(A) (green line, Figure 3A), suggesting that ectopic expression of polyadenylated H3.1 mRNA compromises the assembly of H3.3 nucleosomes around active promoters. Similar results were observed for DNaseI hypersensitive site (DHS)-defined enhancers (Figure 3B) and CTCF binding site-defined insulator regions (Figure 3C). The same results were obtained when the ChIP-seqs were repeated (Figure S2). Collectively, these results indicate that increased H3.1 due to polyadenylation of H3.1 mRNA competes directly or indirectly with H3.3 for deposition in the promoter regions of active genes and other regulatory regions, thereby disrupting proper assembly of H3.3.

**Figure 3.**
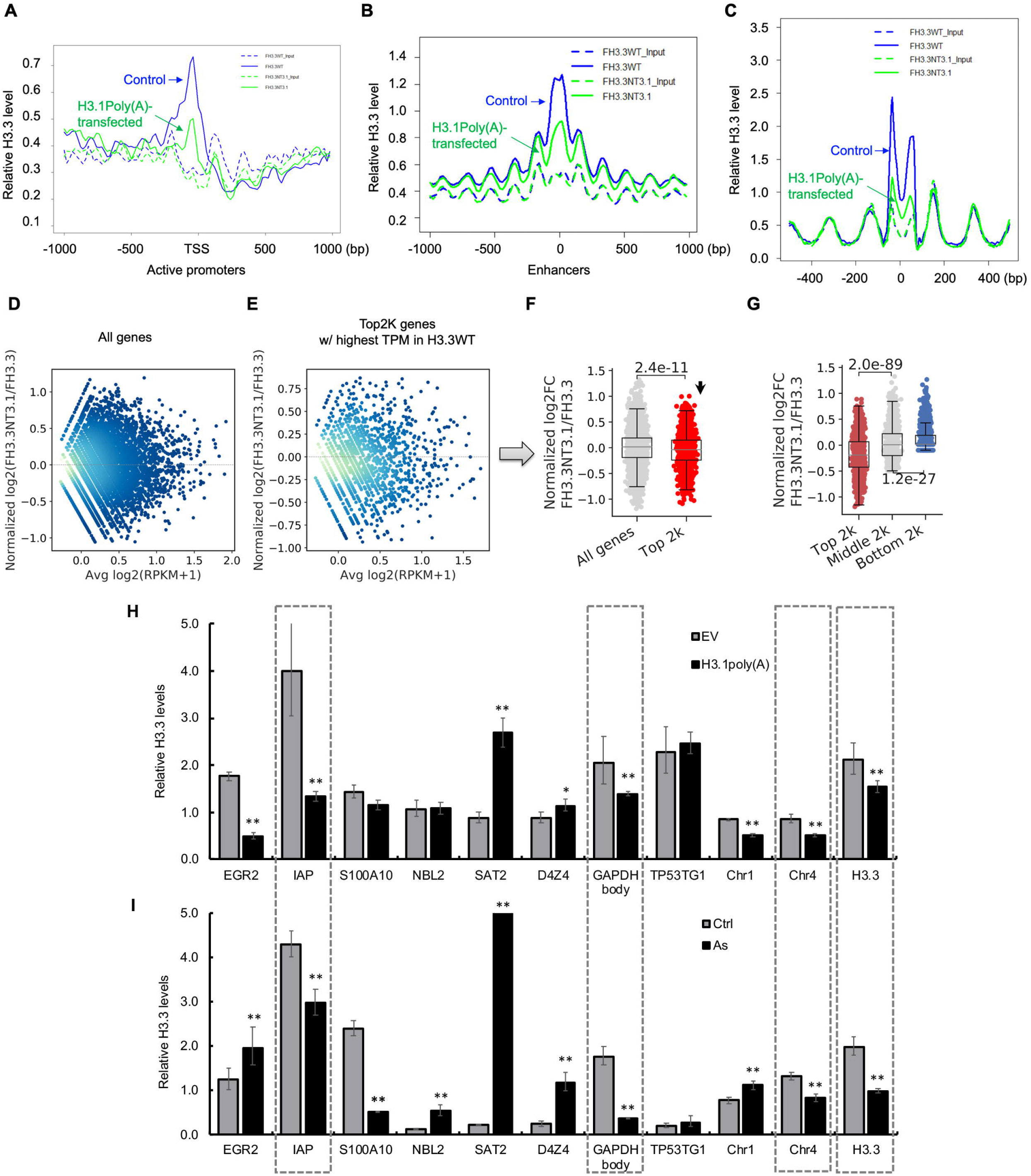
Overexpression of polyadenylated H3.1 mRNA displaces histone variant H3.3 from critical gene regulatory elements. **(A-C)** Profile of H3.3-containg nucleosomes across the transcription start sites (TSSs) for 2,000 highly active genes (activate promoters) (A), DNase I hypersensitive sites (enhancers) (B), or CTCF-binding sites (insulators) (C) are shown. H3.3-containing nucleosomes were isolated followed by ChIP-seq. **(D)** MA plot showing H3.3 level differences in FH3.3NT3.1 compared with FH3.3 on all genes. **(E)** MA plot showing H3.3 level differences in FH3.3NT3.1 compared with FH3.3 on the top 2,000 most highly expressed genes with highest TPM in H3.3 wide type. **(F)** H3.3 level differences of all genes (left) and of the top 2,000 most highly expressed genes (right) in FH3.3NT3.1 compared with FH3.3. P-value was calculated by t-test. **(G)** H3.3 level differences in FH3.3NT3.1 compared with FH3.3 at top (left), middle (middle) and bottom (right) 2,000 genes, ranked by H3.3 level in FH3.3. P-value was calculated by t-test. **(H and I)** The enrichment of H3.3 was measured by ChIP-qPCR at 11 genomic loci in the cells transfected with empty vector (EV) or H3.1poly(A), or treated without (Ctrl) or with arsenic (As). The data shown are presented as mean ±S.D. from qPCRs performed in triplicate. Relative-fold changes were calculated after normalization to input. * *p*<0.05; ** *p*<0.01.

The ChIP-seq profile for H3.3 at active promoters (Figure 3A) measures combined read count (RPKM) of the top 2,000 most highly expressed genes. It does not reflect the changes in each promoter. To explore how H3.3 level in the promoter of each of the top 2,000 highly expressed genes changes by transfection of H3.1poly(A), we calculated the normalized read count (RPKM) around TSS (ranging from −150 bp to +50 bp) of every gene, including the top 2,000 highly expressed genes in FLAG-H3.3 (FH3.3) cells transfected either with untagged H3.1(NTH3.1) or empty vector. We then compared the H3.3 binding difference in FH3.3-NTH3.1 cells vs. that of the FH3.3-Empty cells for each gene (Figure 3D). Although we found unexpectedly, that a large number of genes among the top 2,000 genes displayed increased level of H3.3 after transfection of H3.1poly(A) (Figure 3E), the overall H3.3 level in the promoters of the top 2,000 genes in FH3.3-NTH3.1 cells demonstrated significant decrease compared with FH3.3-Empty cells, in the context of all genes (*p* < 2.4e-11, Figure 3F). These results indicate that H3.3 assembly into the promoters of most highly active genes is more likely to be compromised by polyadenylation of H3.1 mRNA.

To further understand the nature of H3.3 reduction in the promoter regions after transfection of H3.1poly(A), we sorted all genes based on the level of H3.3 enrichment, before the transfection, around TSS (from −150 bp to +50 bp) from strongest to weakest, and then chose the top 2,000, middle 2,000, and bottom 2,000 genes from the list. Next, we compared the extent of H3.3 reduction induced by polyadenylation of H3.1 mRNA among these three sets of genes. While average H3.3 levels in the promoter regions of the middle 2,000 genes were not changed by H3.1poly(A) transfection, the levels were significantly decreased for the top 2,000 genes (Figure 3G). Altogether, these data suggest that genes with high level of H3.3 in their promoters and/or with high expression levels are likely the preferred targets of polyadenylated H3.1 mRNA and susceptible to the resulting increased H3.1 protein and subsequent displacement of H3.3.

Arsenic exposure induces polyadenylation of H3.1 mRNA. To correlate H3.1 mRNA polyadenylation with arsenic effects, we compared the relevance of arsenic exposure and polyadenylation of H3.1 mRNA on H3.3 nucleosome assembly at several genomic loci by ChIP-qPCR. Among 11 locations we tested, H3.3 incorporation decreased at 6 genomic loci by polyadenylated H3.1 mRNA (Figure 3H). H3.3 reduction was also seen after arsenic treatment at 4 out of 6 loci, including IAP (inhibitor of apoptosis) promoter, GAPDH gene body, Chr4 region (α-satellite)[22], and H3.3 promoter (Figure 3I), highlighting the overlapping effects of arsenic exposure and H3.1 mRNA polyadenylation on H3.3 deposition. Given the importance of H3.3 in gene regulation, cell memory, and maintenance of genome integrity, these data indicate that the displacement of H3.3 at important genomic loci through polyadenylation of canonical H3.1 mRNA may be a significant mechanism for arsenic-induced carcinogenesis.

### Ectopic expression of polyadenylated H3.1 mRNA deregulates cancer-associated genes

Reduction of the occupancy of histone variant H3.3 in the critical gene regulatory elements including active promoters, enhancers, and insulator regions by overexpression of polyadenylated H3.1 mRNA (Figure 3), implicates that polyadenylation of H3.1 mRNA might change gene expression. To investigate global changes in transcription induced by ectopic expression of polyadenylated H3.1 mRNA, we carried out RNA-Seqs using BEAS-2B cells stably transfected with untagged-H3.1poly(A) or empty vector. With cutoffs at <0.5 (log_2_FC<-1) and >2 (log_2_FC>1) fold-change (FC) (p<0.01), we identified total 2,597 differentially expressed genes (DEGs) by comparing transcriptomes in H3.1poly(A)-transfected cells and in empty vector-transfected cells. 1,160 and 1,437 genes were found significantly up-or down-regulated, respectively. Next, we performed ingenuity pathway analyses (IPA) on differentially expressed 2,597 genes to characterize cellular pathways associated with the polyadenylation of H3.1 mRNA. Among top 5 diseases and disorders associated with the DEGs, cancer was ranked number 1 (Figure 4A). 467 out of 2,597 H3.1 poly(A)-regulated genes are associated with lung cancer (not shown). We further conducted cancer pathway analysis based on a combined list of DEGs. As shown in Figure 4B, identified key cancer pathways included pancreatic adenocarcinoma, bladder cancer signaling, as well as non-small cell and small-cell lung cancer signaling, analogous to arsenic-targeted cancers [23–25].

**Figure 4.**
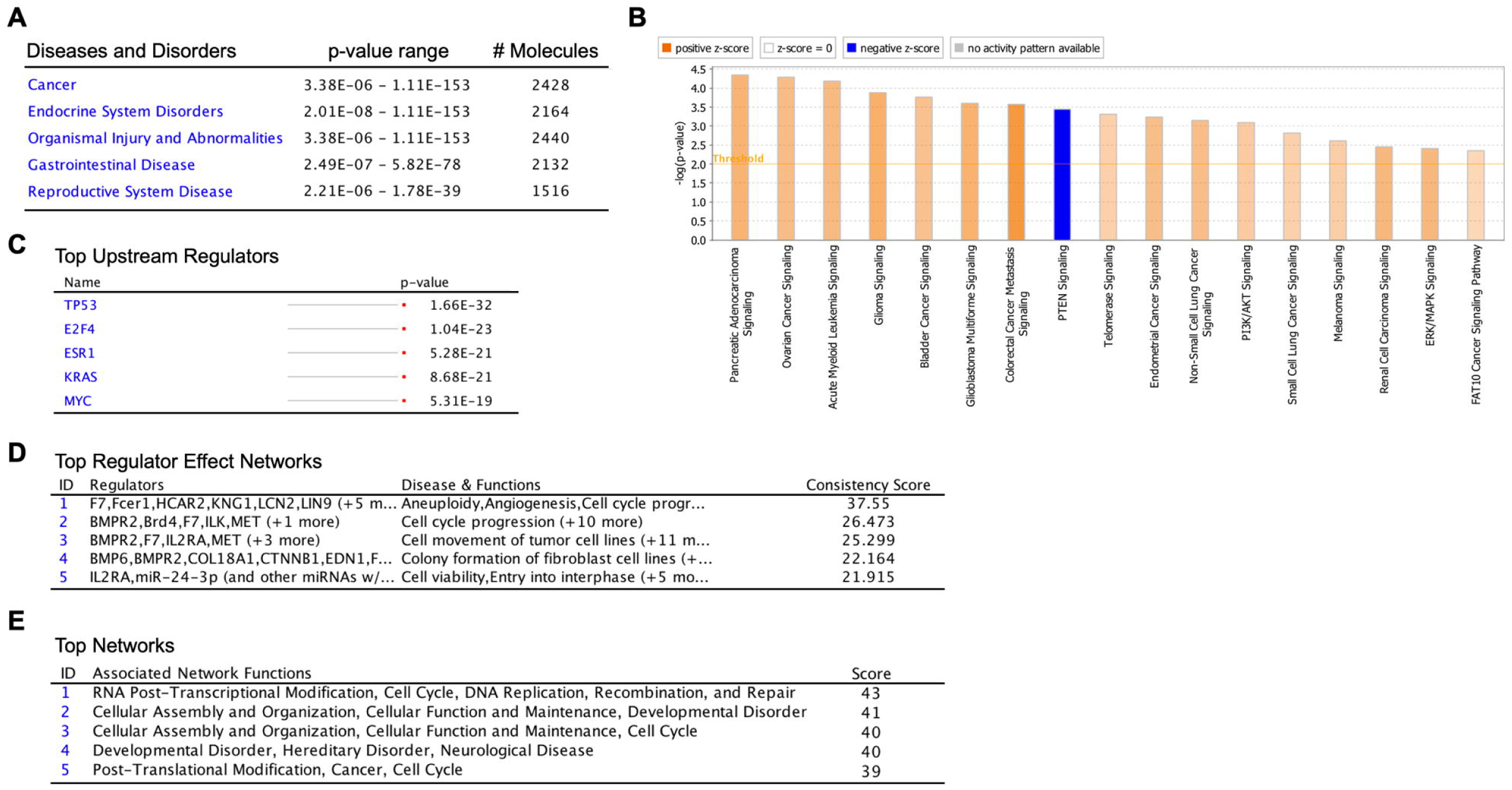
Ectopic expression of polyadenylated H3.1 mRNA deregulates cancer-associated genes. **(A-E)** Ingenuity pathway analyses (IPA) using transcriptome data generated from RNA sequencing (RNA-Seq) of BEAS-2B cells stably transfected with untagged H3.1poly(A) or empty vector. **(A)** Illustrations of top diseases and disorders associated with 2,597 DEGs. **(B)** Illustration of cancer pathway analysis based on 2,597 DEGs. **(C)** Illustrations of top upstream regulators associated with 1,160 up-regulated genes. **(D)** Illustration of top regulator effect network based on combined up-regulated genes. **(E)** Illustration of top networks associated with up-regulated genes.

Interestingly, top upstream regulators of H3.1 poly(A) up-regulated genes included *TP53*, *E2F4*, *ESR1*, *KRAS*, and *Myc* (Figure 4C). High frequency of mutations in *TP53* and *KRAS* genes are common in lung cancers [26], suggesting that H3.1 poly(A) may play roles in lung carcinogenesis through similar pathways initiated by *TP53* and *KRAS* mutations. In addition, analysis of top regulator effect networks of 1,160 up-regulated genes predicted alteration of key cancer-related processes including aneuploidy, angiogenesis, cell cycle progress, cell transformation, and growth of tumor etc. (Figure 4D and Figure S3). Top associated networks for up-regulated genes included RNA post-transcriptional modification, cell cycle, DNA replication, recombination, and repair (Figure 4E). However, top associated networks for down-regulated genes did not include RNA post-transcriptional modification, although cancer was the top disease and one of top networks associated with H3.1 poly(A) down-regulated genes (Figure S4). Together, these transcriptomic analyses suggest that ectopic expression of H3.1 poly(A) can engender extensive differential expression of cancer-related genes, thereby eliciting pathways involved in driving tumorigenic characteristics of human bronchial epithelial cells.

### Polyadenylation of H3.1 mRNA causes mitotic arrest and genomic instability

Polyadenylation of H3.1 mRNA induces disruption of H3.3 assembly at gene regulatory regions such as promoters, enhancers, and insulators (Figure 3A-C). H3.3 is also localized at other critical genomic loci, including pericentric heterochromatin regions and telomeres [20, 27]. The deposition of H3.3 at these regions may also be disrupted by polyadenylation of H3.1 mRNA. Moreover, analysis of upstream regulator effects for the differentially expressed genes by polyadenylated H3.1 mRNA identified cell-cycle progression and aneuploidy, among others, as top affected functions (Figure 4). Thus, polyadenylation of canonical histone H3.1 mRNAs may trigger the dysregulation of cell cycle control and induce chromosomal instability. To examine this possibility, we used flow cytometry to compare alterations in cell cycle between control cells transiently transfected with empty vector and cells transiently transfected with H3.1poly(A) or with H3.1Loop. Significantly less cells were found in G1 phase when transfected with H3.1poly(A) compared to the control and H3.1Loop cells (Figure 5A). Furthermore, cells transfected with H3.1poly(A) experienced a greater G2/M arrest than control and H3.1Loop-expressing cells (Figure 5A). To further distinguish between G2 and M arrest, we used flow cytometry to determine the cellular level of histone H3S10 phosphorylation, a mitosis specific modification. The phosphorylation of H3S10 increased 2-fold upon transfection of H3.1poly(A) plasmid compared with the control cells (Figure 5B). Remarkably, the H3.1Loop-transfected cells (Figure 2A) displayed phosphorylation of H3S10 to the level found with the empty vector cells (Figure 5B). These results suggest that ectopic expression of H3.1 poly(A) mRNA causes mitotic arrest and polyadenylation of its 3’ end is critical for this effect.

**Figure 5.**
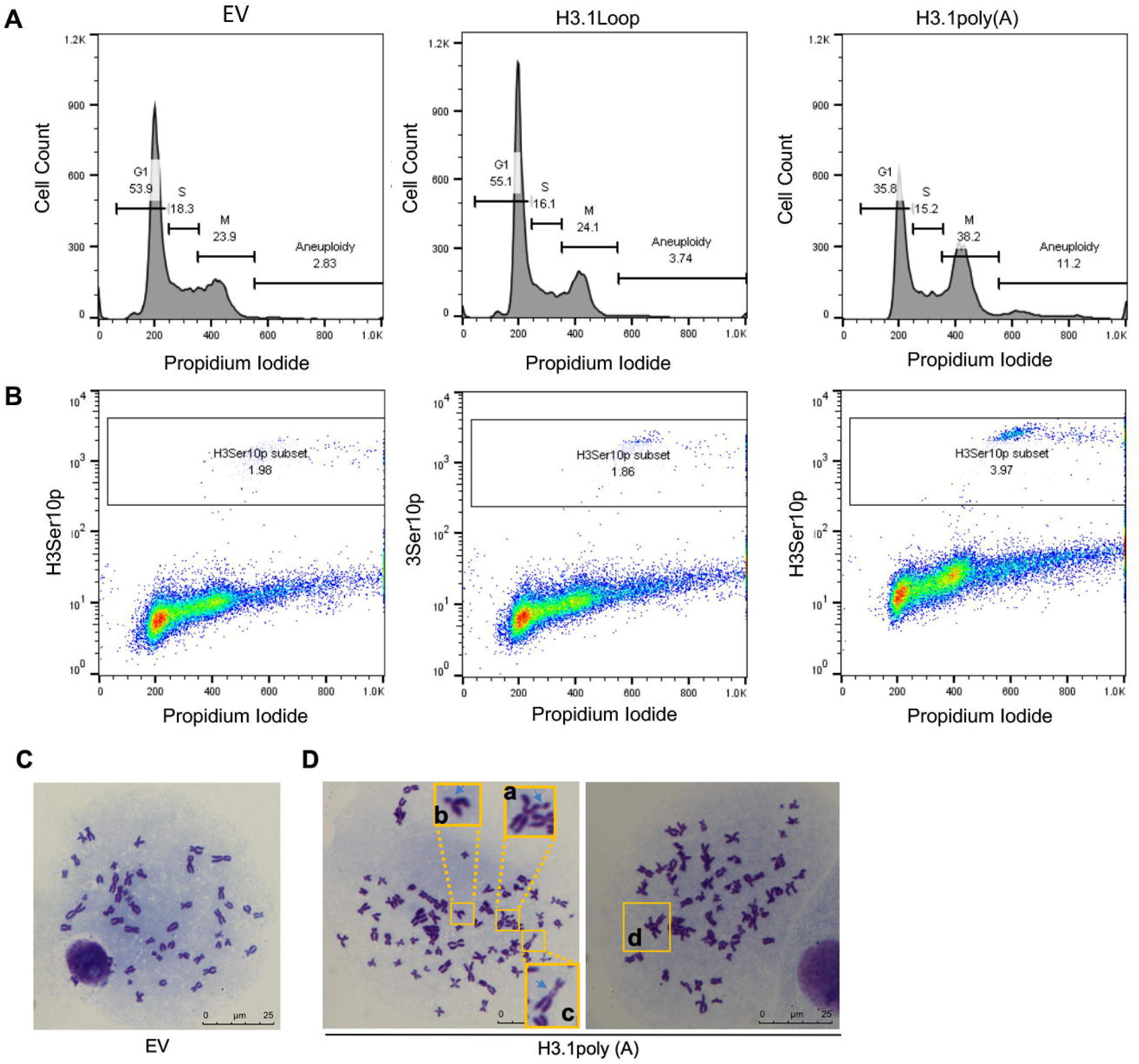
Polyadenylation of H3.1 mRNA causes mitotic arrest and genomic instability. **(A)** Flow cytometry analysis of the cell cycle. BEAS-2B cells that transiently transfected with empty vector (EV), H3.1Loop or H3.1poly(A) were stained with PI. **(B)** Flow cytometry analysis of the cells stained with antibodies specially against phosphorylated H3S10 (H3S10p) to monitor M phase cells. **(C-D)** Giemsa-stained spread chromosomes from BEAS-2B cells transfected with EV or H3.1poly(A) plasmid. (C) Metaphase spread from control cells showing normal chromosome spread. (D) Metaphase spread from H3.1poly(A) cells showing abnormal chromosome spread, including chromatid break (a), deletion (b), dicentric chromosome (c), triradial chromosome (d) etc.

We next examined the possible effects of H3.1 mRNA polyadenylation on chromosome instability. Flow cytometry analysis showed that the number of aneuploid cells increased in H3.1poly(A)-overexpressing cells by almost four-and three-folds when compared with the control cells (from 2.83% to 11.2%) and the H3.1Loop-expressing cells (from 3.74% to 11.2%), respectively (Figure 5A), indicating that polyadenylation of H3.1 mRNAs causes defective chromosome segregation. To identify chromosomal aberrations induced by polyadenylated H3.1 mRNA, we performed cytogenetic analysis using BEAS-2B cells transfected with either empty vector or H3.1poly(A) plasmid. While empty vector-transfected cells displayed normal metaphase spread (Figure 5C), abnormal chromosomes such as deletions, chromatid breaks, dicentric and triradial chromosomes were observed from H3.1poly(A)-transfected cells (Figure 5D). Collectively, these results suggest that polyadenylation of H3.1 mRNA can induce mitotic arrest and genomic instability.

### Ectopic expression of H3.3 attenuates arsenic-induced anchorage-independent cell growth

Polyadenylation of H3.1 mRNA induced by arsenic exposure compromises assembly of H3.3 at important genomic loci. Numerous findings linked abnormal H3.3 to cancer initiation and development [1, 20, 28–31]. Moreover, H3.3-knockout mice were embryonically lethal and displayed severe chromosomal aberrations [31, 32], suggesting that disruption of H3.3 assembly might be a significant contributor to arsenic-induced carcinogenesis. To test this hypothesis, we first examined the effect of H3.3 knockdown on anchorage-independent cell growth. Both H3.3 mRNA levels (Figure 6A) as well as protein levels (Figure 6B) were reduced by transfection of two different types of H3.3 siRNAs in BEAS-2B cells as compared with transfection of the control siRNA. Soft agar assays showed that knocking down histone H3.3 expression enhanced soft agar colony formation of BEAS-2B cells (Figure 6C). To further examine whether inhibition of H3.3 assembly was the underlying defect in arsenic-induced cell transformation, we overexpressed H3.3 (Figure 6D), which could antagonize arsenic-induced disruption of H3.3 assembly by increasing histone H3.3 supply, followed by soft agar assays. Anchorage-independent cell growth was facilitated by treatment of BEAS-2B cells with arsenic either for 3 days at 1 μM or 1 week at 0.5 μM (Figure 6E). Overexpression of H3.3 attenuated arsenic-induced colony formation on soft agar (Figure 6E), suggesting that aberrant histone H3.3 assembly was a significant contribution to arsenic-induced cell transformation.

**Figure 6.**
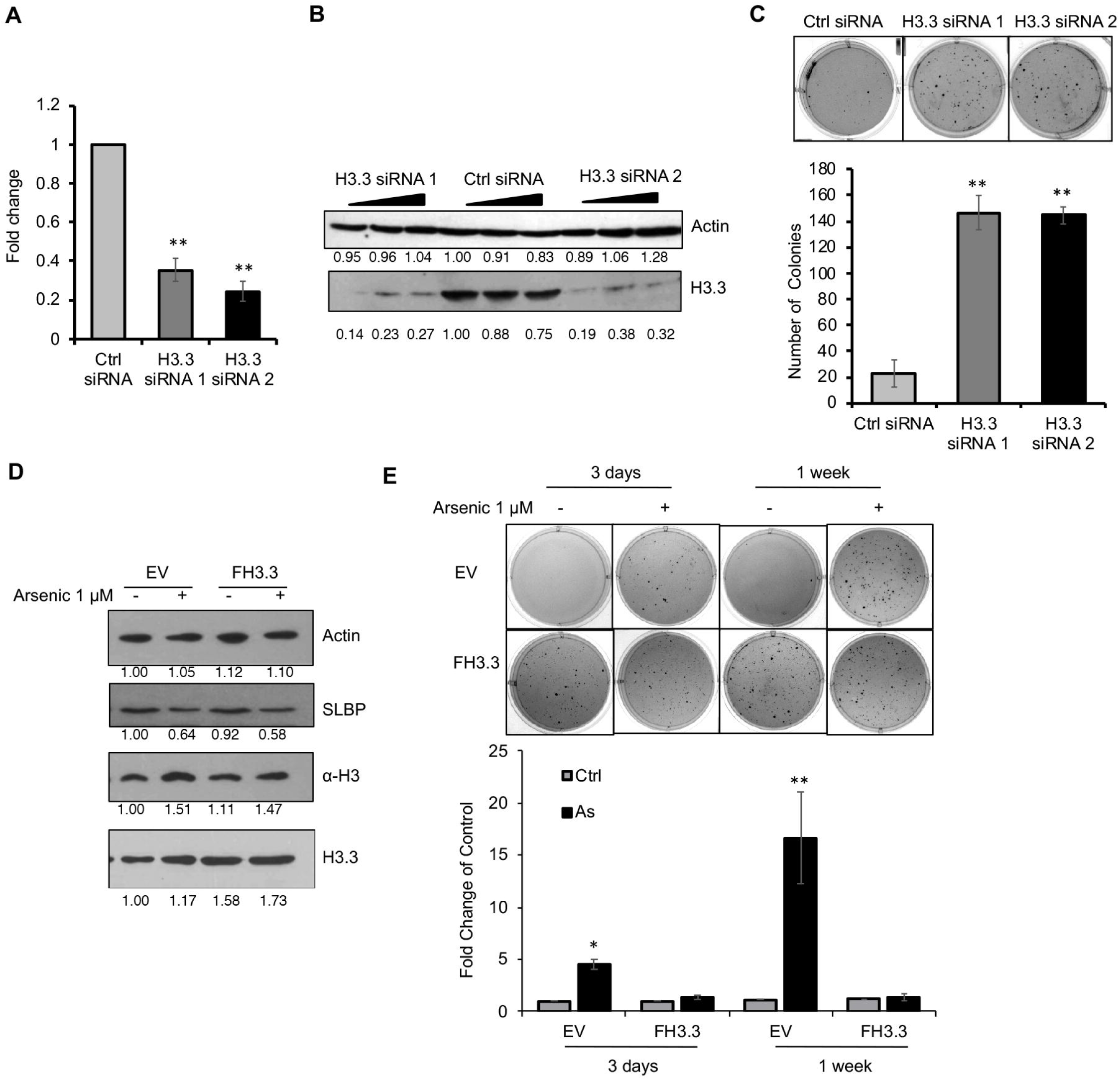
Ectopic expression of H3.3 attenuates arsenic-induced anchorage-independent cell growth. **(A-C)** Knockdown of H3.3 facilitates colony formation of BEAS-2B cells on soft agar. **(A)** RT-qPCR analysis of *H3.3* mRNA levels in BEAS-2B cells after 48-h transfection with control (Ctrl) siRNA or with two distinct siRNAs for H3.3. **(B)** Western blot using indicated antibodies. **(C)** Soft agar assays. The data shown are the mean ±S.D.(n=3). ** *p*<0.01. **(D-E)** Overexpression of H3.3 rescues arsenic-induced cell transformation. BEAS-2B cells were transiently transfected with pcDNA-empty (EV) or with pcDNA-FLAG-H3.3 (FH3.3) in the presence or absence of 1 μM arsenic. **(D)** Western blot using indicated antibodies. **(E)** Soft agar assays. The data shown are the mean ±S.D. (n=3). * *p*<0.05; ** *p*<0.01.

Polyadenylation of H3.1 mRNA/inhibition of H3.3 assembly appeared critical for arsenic-induced cell transformation. To further assess the importance of this pathway in arsenic carcinogenesis, we tested whether overexpression of SLBP, which should compensate for the arsenic-mediated depletion of SLBP thereby inhibiting generation of polyadenylated H3.1 mRNA, rescues arsenic-induced cell transformation. BEAS-2B cells were stably transfected either with empty vector or SLBP-expressing plasmid. Western blot and RT-qPCR results showed that the protein and mRNA levels of SLBP were increased by about 3-fold in the SLBP overexpressing clone as compared with the empty vector control (Figure S7A and B). Whereas the level of polyadenylated H3.1 mRNA increased after arsenic treatment in control cells, arsenic-induced polyadenylation of H3.1 mRNA was significantly attenuated in the cells that stably express SLBP (Figure S7C). Similarly, overexpression of SLBP apparently inhibited arsenic-induced anchorage-independent growth of BEAS-2B cells (Figure S7D), indicating an essential role for SLBP depletion/H3.1 mRNA polyadenylation/disruption of H3.3 assembly in arsenic-induced cell transformation.

### Gain of polyadenylated H3.1 mRNA by arsenic exposure *in vivo*

To determine whether arsenic is also able to reduce SLBP and alter canonical histone H3.1 mRNA processing *in vivo,* we measured levels of SLBP and polyadenylation of H3.1 mRNA in male A/J mice exposed to arsenic (0, 100, and 200 μg/L) by inhalation for one week. Lung tissues were collected from the mice and RT-qPCR was first employed to study H3.1 polyadenylation and levels of SLBP mRNA after arsenic exposure. RT-qPCR results showed that polyadenylated H3.1 mRNA levels were elevated with increasing arsenic exposure (Figure 7A). A reduction in SLBP mRNA was observed most significantly in the 200 μg/L treatment group (Figure 7B). Western blot analysis further demonstrated a downward trend in SLBP protein levels upon arsenic exposure (Figure 7C). SLBP is essential for the processing of canonical histone pre-mRNA, thus a loss of SLBP would cause polyadenylation of canonical histone mRNAs *in vivo*. The notion that loss of SLBP and gain of polyadenylated H3.1 mRNA occur *in vivo* suggests that polyadenylation of canonical histone H3.1 mRNA may play roles in arsenic-induced toxicity and carcinogenesis.

**Figure 7.**
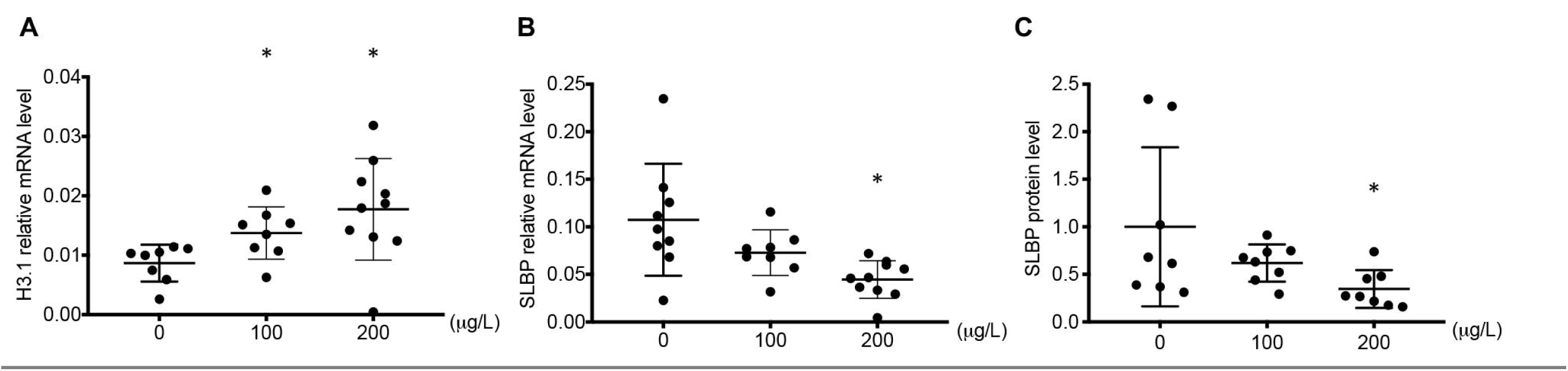
Gain of polyadenylated H3.1 mRNA and loss of SLBP by arsenic exposure *in vivo*. **(A-C)** A/J mice were treated with NaAsO_2_ for 1 week. Lung tissues were extracted and mRNA or protein levels were analyzed by RT-qPCR (A,B) or Western blot (C). The data shown for each individual mouse. Relative protein levels were calculated based on band intensity. Error bars represent S.D. * *p*<0.05.

## Discussion

We previously found that arsenic exposure induces polyadenylation of canonical histone H3.1 mRNA [9]. Our present study demonstrates that polyadenylated H3.1 mRNA induces chromosome instability and cell transformation likely by displacing its specific replacement histone H3.3 from critical genomic loci in mammals.

What is the possible mechanism(s) by which overproduced H3.1 protein by polyadenylated H3.1 disrupts H3.3 assembly? The polyadenylation of H3.1 mRNA elevates cellular H3.1 protein, resulting in excess H3.1 not only during S phase but also outside of S phase [9]. While H3.1 is incorporated into the chromatin exclusively by the replication-dependent chromatin assembly factor 1 (CAF-1) complex, many histone chaperones such as HIRA, DAXX, and DEK, have been identified as H3.3-specific histone chaperones for replication-independent H3.3 assembly [1, 20, 33]. CAF-1 complex is composed of three subunits, i.e., p150, p60, and p48. Whereas p150 is specific for H3.1, p60 and p48 exist in HIRA complex as well [16]. Thus, excess H3.1 protein might increase the association of H3.1 with CAF-1, including histone chaperones p60 and p48, which will result in a shortage of these two chaperones available for HIRA and in turn decrease the interaction between H3.3 and the HIRA complex, thereby compromising H3.3 assembly. Another scenario would be that excess H3.1 protein during M phase may directly compete with H3.3 variant for deposition onto genomic loci with higher H3.3 turnover. This appears to be possible since the overexpression of canonical H3.1 increased replication-independent H3.1 assembly outside of S phase in flies [34]. Moreover, a study showed that altering the ratio of H3.1 to H3.3 by increasing H3.1 protein resulted in replacement of H3.3 with H3.1 in regulatory regions of skeletal muscle cells [35]. Inhibition of H3.3 assembly did not appear to result from downregulation of H3.3-specific chaperones based on our RNA-seq data (not shown). However, we cannot rule out the possibility that post-translational modifications of H3.3 could be altered indirectly by H3.1 polyadenylation, contributing to altered assembly of H3.3 nucleosomes. The reduction of H3.3 at these genomic loci was not due to the downregulation of FLAG-H3.3 expression, since (a) FLAG-tagged H3.3 protein levels were not decreased after transfection of H3.1poly(A) plasmid (Figure S1), and (b) FLAG-H3.3 levels were not decreased at all the genomic loci we tested, but in fact, were slightly increased in the promoters of most silenced genes (Figure 3G). In addition, while H3.3 displacement was more likely to occur in the promoters of highly expressed genes as well as genes whose promoters are enriched with H3.3 (Figure 3F and G), we did not observe a direct relationship between the extent of H3.3 loss and gene downregulation (not shown). Therefore, the H3.3 reduction was not likely a secondary event following transcriptional downregulation by H3.1 polyadenylation.

Displacement of H3.3 from critical genomic loci could have significant impact on carcinogenesis. H3.3 depletion resulted in cell death and mitotic defects, leading to karyotypical abnormalities such as lagging chromosomes, aneuploidy, and polyploidy [31]. H3.3-knockout mice exhibited early embryonic lethality [31, 32]. Moreover, recent studies have directly linked histone variant H3.3 to oncogenesis. Sequencing studies revealed a high frequency of mutations in H3.3 genes in several cancer types including pediatric high-grade glioblastoma and bone tumors [36–39]. Downregulation of H3.3 expression in adult glioblastoma has been found to disrupt chromatin organization and promote cancer stem cell properties [40]. Besides mutation and transcriptional downregulation, disruption of the H3.3 assembly has also implicated in cancers [38, 41–45]. For example, somatic mutations in the DAXX and ATRX, the histone chaperones for H3.3 deposition at telomeric and pericentromeric regions, were identified in pediatric glioblastoma tumors [46–49]. 43% of pancreatic neuroendocrine tumors (PanNETs) had mutations in genes that encode ATRX and DAXX [44]. A DAXX missense mutation was also found in acute myeloid leukemia [50]. Mutation of DEK significantly reduced H3.3 loading in some human myeloid leukemia patients [51, 52]. These studies indicate an association between disruption of H3.3 assembly with cancer causation. In the present study, we demonstrate that arsenic-induced polyadenylation of canonical H3.1 mRNA leads to inhibition of H3.3 assembly at critical genomic loci by disrupting the stoichiometry of replication-dependent and-independent histones. Our findings thus add new insight into mechanisms by which histone variant H3.3 exerts a role in tumorigenesis.

Appropriate histone stoichiometry is essential for maintaining genomic integrity. Histone stoichiometry could be altered by either decreased or increased protein levels for a particular histone gene. While it has been relatively well known that decreases in histone protein level can induce genomic instability by facilitating chromatin accessibility [53–56], the mechanisms that underlie genomic instability induced by the histone increase are poorly understood [57, 58]. Our present study demonstrates that upregulation of canonical histone protein may induce chromosome instability by displacing their specific replacement histones from critical genomic loci in mammals.

In *Drosophila*, SLBP depletion led to polyadenylation of all core histones [5, 6, 8], implicating that mRNAs for other canonical histones besides H3.1 may also be polyadenylated by arsenic exposure. In fact, RT-qPCR results show that the levels of polyadenylated mRNAs for H2A, H2B, H3.2, and H4 were increased after arsenic exposure (Figure S5A-E). The protein levels for H2A, H2B, total H3, and H4 were also upregulated by arsenic treatment (Figure S6A-E). The increase in canonical histone proteins other than H3.1 may also affect chromatin assembly and structure. However, we anticipate that the impact of polyadenylation of canonical histone H3 mRNAs might be the greatest. H3.3 seems to be the only histone variant that serves as a “replacement histone”. For example, over the lifespan of the mouse, H3.3 replaces canonical H3 and becomes the predominant H3 protein in non-dividing cells. Although histone H2A has several variants, such as H2A.X, H2A.Z, and macroH2A, they are not likely serving as “replacement histones”. Based on elegant work from Marzluff’s group, a subset of canonical histone genes are expressed independently of DNA replication as polyadenylated mRNAs, and likely serve as “replacement histones” for the H2A, H2B, and H4 in terminally differentiated cells [59]. The implication of these observations is that while mRNAs for all canonical histone genes are polyadenylated after arsenic treatment and produce more respective proteins, increased canonical H3 proteins, i.e., H3.1 and H3.2, may have a greater impact on chromatin structure since they can displace the “replacement histone” H3.3, which has distinct features from canonical H3.

We previously reported a marked increase in polyadenylated mRNA for H3.1 in arsenic-transformed BEAS-2B clones, but not in chromium (VI)-transformed BEAS-2B [9]. Depletion of SLBP and increase in polyadenylated H3.1 mRNA was also observed following nickel treatment, whereas polyadenylation of H3.1 mRNA was not evident in cadmium-exposed cells [60]. These data suggest that the polyadenylation of canonical histone mRNAs is specific to certain metals. Since both arsenic and chromium (VI) induce oxidative stress, it seems unlikely that oxidative stress per se is the cause of polyadenylation of canonical histone mRNAs. SLBP depletion appears to be a key factor for arsenic-induced polyadenylation of canonical histones, since the transcription of other genes involved in canonical histone mRNA processing, including U7 snRNP (LSM10), CPSF2, 3, XRN2, and ZNF473, was not changed by arsenic exposure [9]. Arsenic and nickel-induced SLBP depletion is epigenetically regulated by its promoter and also by proteasomal degradation. It would be important to identify which histone/DNA modifying enzymes or protein kinases/phosphatases are involved in regulating arsenic-induced SLBP depletion [61–63]. Arsenic is able to release zinc from proteins that contain 3-cysteine or 4-cysteine zinc finger motif thereby changing protein structures and enzyme activities [64], whereas nickel inhibits enzymatic activities of dioxygenases such as histone demethylase JMJD1A by replacing the ferrous iron in the catalytic centers [65]. Therefore, it would be interesting to examine if the above-characterized enzymes contain zinc finger motif or require iron for their activity.

In summary, we demonstrated that arsenic-induced polyadenylation of canonical histone H3.1 mRNA is able to displace H3.3 variant from promoters, enhancers, and insulator regions, resulting in transcriptional deregulation, cell cycle arrest, genomic instability, and tumor formation in nude mice. We uncovered polyadenylation of canonical histone H3.1 and disruption of H3.3 assembly as a potential new mechanism for arsenic-induced carcinogenesis. This sheds new light on the oncogenic role for H3.3. Our findings also provided a new perspective on genomic instability induced by alterations of histone stoichiometry. In the future, we will address to what extent polyadenylation of canonical histones accounts for the effects of arsenic, and by what mechanisms arsenic induces SLBP depletion and increased H3.1 displaces H3.3 at regulatory regions. The findings from these studies will provide us potential therapeutic and preventive targets for arsenic-induced cancers.

## Acknowledgements

We thank the NYULMC Genome Technology Center, partially supported by the Cancer Center Support Grant P30CA016087. The authors thank Thomas Des Marais and Gabriele Grunig for technical assistance and Suresh Cuddapah for helpful comments. This work was supported by grants from the US National Institutes of Health: R01ES026138 (M.C. and C.J.), P30ES000260 (M.C. and Pilot Project Program to C.J.), R01ES029359 (M.C. and C.J.), R01ES022935 (M.C.), and K22CA204439 (C.Z.).

## Author Contributions

C.J and M.C conceived the study. D.C. and Q.Y.C. performed most of the experiments and data analysis. Z.W. performed most of the computational analysis under supervision of C.Z.. Y.Z., T.K., W.T., J.L., F.W., L.F., X.Z., R.H., S.S., and H.S. performed experiments and/or data analysis; Q.Y.C., D.C., Z.W., C.Z., C.J., and M. C. wrote the manuscript.

## Declaration of Interests

The authors declare no competing interests.

## STAR Methods

### Cell Lines

Immortalized human bronchial epithelial (BEAS-2B) cells were obtained from the American Type Culture Collection (ATCC, Manassas, VA). BEAS-2B cells were adapted to serum growth immediately after their purchase and have been carefully maintained at below confluent cell density. The cells were recently tested through Genetica DNA Laboratories and proved to be 100% authentic against a reference BEAS-2B cell line (Burlington, NC). Cells are cultured in Dulbecco’s Modified Eagle Medium (DMEM, Invitrogen, Grand Island, NY) supplemented with 1% penicillin/streptomycin (GIBCO, Grand Island, NY) and 10% heat-inactivated fetal bovine serum (FBS, Atlanta Biologicals, Lawrenceville, GA). All cells were cultured in a 37°C incubator containing 5% CO_2_. BEAS-2B cells were authenticated by Genetica DNA Laboratories (Burlington, NC) on July 22, 2015. Cells were matched 100% to 15 short tandem repeat (STR) loci and amelogenin to the reference profile of BEAS-2B (ATCC CRL-9609). For arsenic exposure, cells were treated with sodium meta-arsenite (NaAsO_2_, Sigma, St. Louis, MO) with doses ranging from 0 to 2 μM for 0 to 96 hr.

### Animals, arsenic exposure, and sample collection

7-week-old male A/J mice (SPF grade) were purchased from Jackson Laboratories (Bar Harbor, ME) and maintained at the New York University School of Medicine animal facility under standard environmental conditions (22 °C, 40-70% humidity, and a 12:12-hour light: dark cycle). All mice were handled in accordance with NIH and Institutional Animal Care and Use Committee (IACUC) guidelines. A purified rodent diet was provided. The mice were observed for one week before the start of the experiment. Sodium arsenite exposure treatment was prepared at 0, 100, and 200 μg/L concentrations, and the mice were given access to the exposure via intratracheal instillation every other day for 1 week. Post-treatment, the mice were sacrificed by CO_2_ and lung tissues were extracted. Protein lysates were extracted using boiling buffer and analyzed for SLBP levels using Western blotting using β-actin as internal control. RNA was extracted using standard Trizol protocol and analyzed for H3.1 poly(A) and SLBP mRNA using qPCR using GAPDH as loading control. All procedures were conducted in compliance with New York University’s guidelines for ethical animal research and the Declaration of Helsinki.

### Xenograft tumor model

18 Athymic NCr-nu/nu (5-6-week-old) mice were obtained from the National Cancer Institute (NCI, Frederick, MD) and housed in a pathogen-free facility for all experiments. A total of 5 million control or H3.1poly(A)-transfected cells were suspended in 0.1 mL PBS and subcutaneously injected into each side of the femoral area of the same mouse. Each mouse is injected on one side with control and the other with H3.1-transformed cells to ensure the same biological environment. Each cell line was subcutaneously injected into three mice, in triplicates. Tumor growth and overall health of the mice were monitored once a week. At 5 months post-injection, the mice were sacrificed by CO_2_ euthanization and the tumors were extracted by standard surgery for determination of tumor weight. Cut-off weight for tumor weight was 0.1g. Tumor size was measured using calipers at the indicated times. The tumor volume was calculated according to the formula: volume = length*(width^2^)/2. Graphs illustrating tumor number, weight, and volume were generated using Prism 7 7.0e (GraphPad Software). Animal experiments in the present study were performed in compliance with the guidelines of The Center for Laboratory Animal Research, New York University School of Medicine.

### Plasmids and Cell Transfection

pcDNA3.1-H3.1-poly(A) and pcDNA3.1-FLAG-H3.1-poly(A) plasmids were previously constructed in our lab. A 110 bp DNA fragment of H3.1 gene containing the stem loop sequence and 5′ and 3′ linker sequences for sub-cloning into the XbaI and ApaI sites were amplified by PCR using the human genomic DNA as a template. The fragment was inserted into the pcDNA3.1-H3.1-poly(A) plasmid to obtain pcDNA3.1-H3.1-Loop plasmids. Transfections were carried out using PolyJet (SignaGen, Rockville, MD) according to the manufacturer’s instructions. pcDNA3.1-empty vector was used as the control. Primers: forward 5′-CTAGTCTAGAGTCTGCCCGTTTCTTCCTC-3′; reverse5′-CCGGGCCCAAACTAACATAGACAACCGA-3′. Control short interfering RNA (siRNA) or two distinct siRNAs for H3.3 (H3.3 siRNA1 and H3.3 siRNA 2) (Sigma, St. Louis, MO) were transfected into BEAS-2B cells to knock down H3.3 expression. Transfections were carried out using LipoJet (SignaGen, Rockville, MD) according to the manufacturer’s instructions.

pcDNA-FLAG-SLBP plasmids were purified using a Qiagen QIAprep Spin Midiprep kit prior to transfection. Overexpression transfections were performed using Lipofectamine® LTX Reagent with PLUS reagent (Invitrogen, Grand Island, NY) following the manufacturer’s protocol. Briefly, 150,000 cells were seeded into 6-well dishes 24 hours prior to transfection. The following day, 1 ug of purified plasmid was transfected into each well using 10 μL of Lipofectamine LTX and 2.5 μL of PLUS reagent per transfection. 24 hours post-transfection, the media was removed and replaced with fresh DMEM. After three days, 0.5 μg/ml of G418 selection agent was added to the transfected cells. The cells were grown under selection for three weeks and harvested for western blot and qPCR analysis.

### Antibodies

| Name | Source | Catalog Number |
| --- | --- | --- |
| Histone H3 | Abcam | ab1791 |
| SLBP | Abcam | ab 181972 |
| β-actin | Abcam | ab8226 |
| Histone H4 | Millipore | 07-108 |
| Histone H2A | Millipore | 07-146 |
| Histone H2B | Millipore | 07-371 |
| FLAG | Sigma | F3165 |

### Western blotting

Cells were lysed with boiling buffer (1% SDS, 10 mM Tris (pH 7.4), 1 mM sodium orthovanadate) and 50 μg of whole cell lysate were separated by 12% SDS-PAGE and transferred to a PVDF membrane. After blocking in 5% skim milk in TBST for 1 h at room temperature, the membrane was incubated with primary antibodies overnight at 4 °C, and then probed with HRP labeled secondary antibody (1:2000) for 1 h at room temperature before the visualization by the chemiluminescence. Quantification of immunodetected proteins was performed using Image J software.

### RNA Isolation and Quantitative Real-time PCR

Total RNA was extracted using TRIzol (Invitrogen, Grand Island, NY) and subsequently synthesized into single-stranded cDNA using ProtoScript® II First Strand cDNA Synthesis Kit (New England BioLabs, Ipswich, MA) in accordance with manufacturer’s instructions. 1μg of RNA in a final volume of 20 μl. Quantitative real time PCR analysis was performed using Power SYBR Green PCR Master Mix (Qiagen) on the ABI PRISM 7900HT system. All experiments were performed in triplicates. Relative gene expression levels were normalized to either Tubulin or GAPDH expression. The results were presented as–fold change to the level expressed in control cells.

### Primers Used

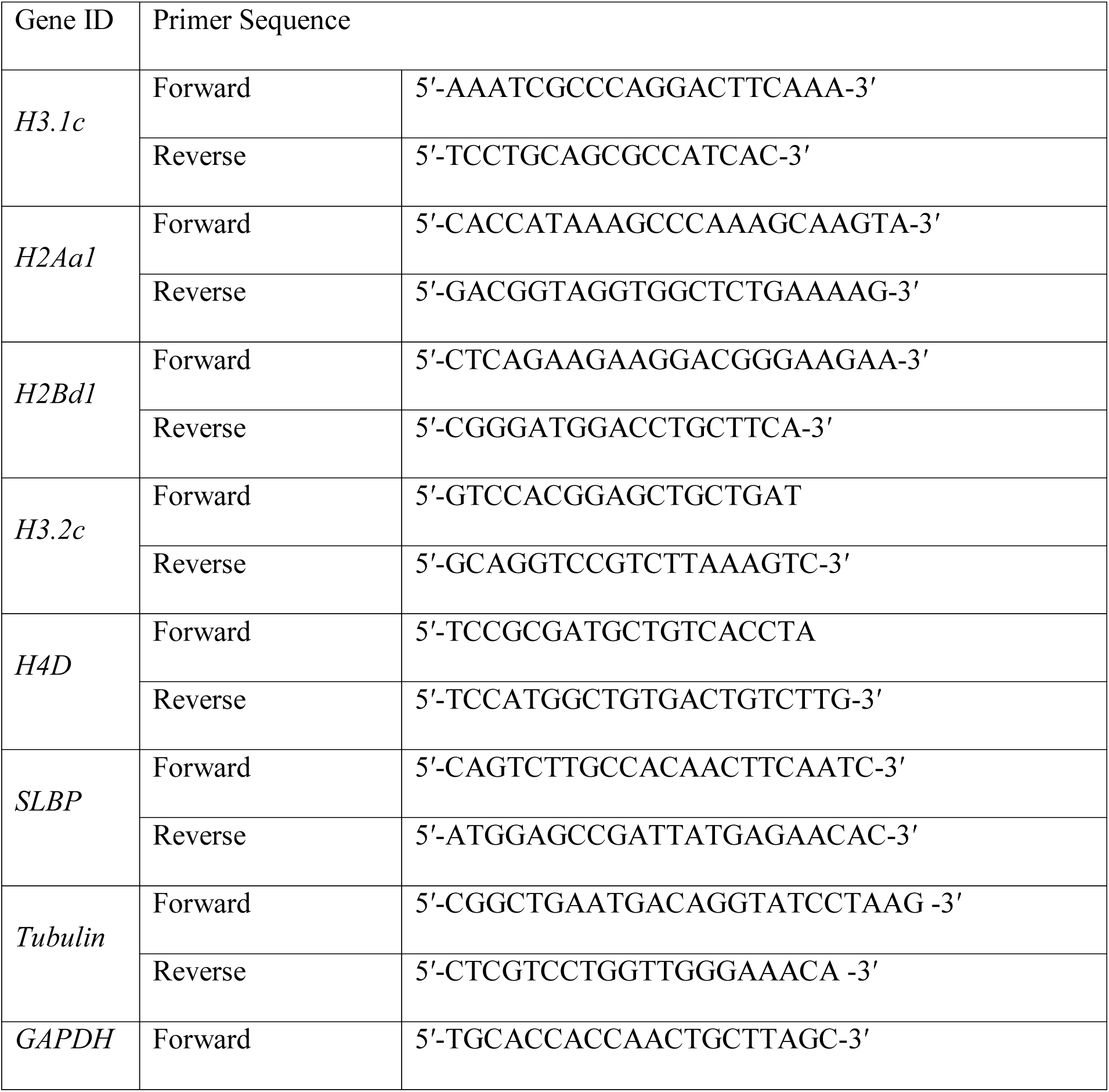
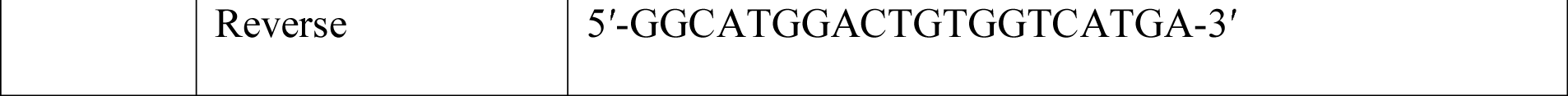

### Reverse Transcriptase-Polymerase Chain Reaction (RT-PCR)

The total RNA of treated cells was extracted using the Trizol reagent (Invitrogen, Grand Island, NY); 1 ug of RNA was reverse transcribed using the SuperScript IV First-Strand Synthesis System (Invitrogen, Grand Island, NY) following the manufacturer’s protocol. Amplification cycles were: 95 °C for 5 min and then 30 cycles at 95 °C for 30 s, 58 °C for 30 s, 72 °C for 30 s, followed by 72 °C for 10 min. Aliquots of PCR products were checked by electrophoresis on a 2% agarose gel with the fragments visualized by ethidium bromide staining.

### Primers Used

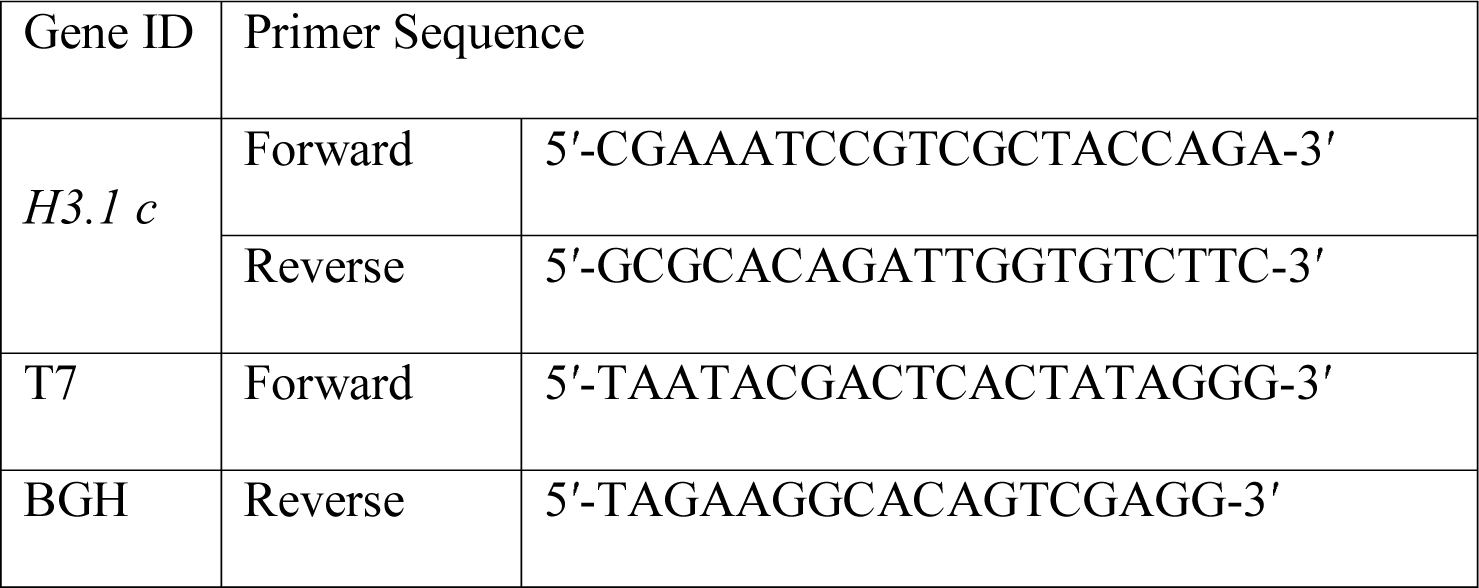

### Cell proliferation assay

Cells were plated in 96-well plates at a density of 5*10^3^ cells per well and incubated at 37 °C in 5% CO_2_. At different time points (0, 1, 2, 3, 4, 5, 6, 7 days), the medium was replaced with 100 μl fresh medium containing 0.5 mg/ml 3-(4,5-dimethylthiazol-2-ul)-2,5-diphenyl tetrasodium bromide (MTT). Four hours after the addition of MTT, 100 μl of isopropanol containing HCL was added to each well to dissolve the crystals. Fluorescence was monitored using a microplate reader (Spectra Max M2, Molecular Devices) at a wavelength of 570 nm. Each experiment was performed in eight times and repeated three times. Data was reported as mean ± SD.

### Anchorage independent growth assay

Cells were rinsed with PBS to remove the metal from the media then seeded in low gelling temperature Agarose Type VII (Sigma Aldrich, St. Louis, MO). 5,000 cells were seeded in triplicate in 6-well plates in a top layer of 0.35% agarose onto a bottom layer of 0.5% agarose. Cells were allowed to grow for four weeks until individual colonies were large enough to be select from the agar. Colonies were picked from each treatment and control group. These colonies were grown out into monolayers for four weeks. After monolayer growth, cells were collected in Trizol for quantitative real-time PCR (RT-qPCR). A second set of plates was stained with INT/BCIP solution (Roche Diagnostics, Indianapolis, IN) for visualization and quantification of colony growth in agar, according to the manufacturer’s protocol.

### Flow Cytometry

For flow cytometry cell cycle analysis, the cells were fixed with 4% formaldehyde for 10 min at RT and permeabilized by adding cell suspension drop-wise into ice-cold 100% methanol while gentle vortexing to a final concentration of 90% methanol and incubate on ice for 30 min. Permeabilized cells were resuspended in FACS buffer (1xPBS, 1% BSA), stained with Alexa-488-conjugated phospho-Histone H3 (ser10) (1:50, CST9708, Cell Signaling Technology, Danvers, MA) for 1 hr at room temperature in dark. Then cells were counterstained with propidium iodide (Sigma-Aldrich, St. Louis, MO) 30 min at room temperature in dark. Stained cells were counted and analyzed via FACS Calibur flow cytometer (BD, Bioscience, San Jose, CA).

### Transwell Cell Invasion assay

For cell invasion analysis, Biocoat Matrigel® Invasion Chambers (Corning, Corning, NY) were incubated with DMEM containing 0.1% FBS at 37°C for 3-h. BEAS-2B cells were trypsinized, centrifuged, and washed to remove excess FBS. 5×10^4^ cells were resuspended with 400 μL of DMEM containing 0.1% FBS. The cells containing DMEM with 0.1% FBS were seeded in wells containing DMEM with 10% FBS and grown for 24-h in a 37°C incubator containing 5% CO_2._ The migrated cells were fixed with 3.5% formaldehyde for 5 minutes at room temperature and subsequently incubated with methanol for 20 minutes. The wells were then incubated with 5% Giemsa solution overnight at room temperature and washed with water the next day for observation under the microscope.

### Chromosome Spread

Transfected cells were arrested in metaphase by the addition of 0.1 μg/ml colcemid for 3 hr. The cells were trypsinized and collected by centrifuge at 1,000 rpm 5 min. Hypotonic treatment was accomplished with 0.075 M KCl at 37°C for 17 min, followed by fixation in 3:1 methanol–acetic acid (three changes). Prepared cells were dropped on cold, wet slides, stained for 5 min in 10% Giemsa (Sigma-Aldrich, St. Louis, MO) in pH 7.0 phosphate buffer and observed under the light microscope with oil lens.

### Nucleosome Preparation, ChIP and qPCR Analysis

Mono-and dinucleosomes were isolated by Micrococcal Nuclease (MNase) digestion and sucrose gradient purification from arsenic-treated and untreated BEAS-2B cells as described (Jin and Felsenfeld 2006). ChIP was performed using anti-FLAG M2 Affinity-gel (A2220, Sigma-Aldrich, St. Louis, MO). DNA fragments were purified by phenol/chloroform extraction and amplified by real-time quantitative PCR in 7900 Fast Sequence Detection System (Applied Biosystems, Foster City, CA). The used primers represent different regions of genome, including pericentric, telomere, heterochromatin regions and transcription starting sites, see below.

### Primers Used

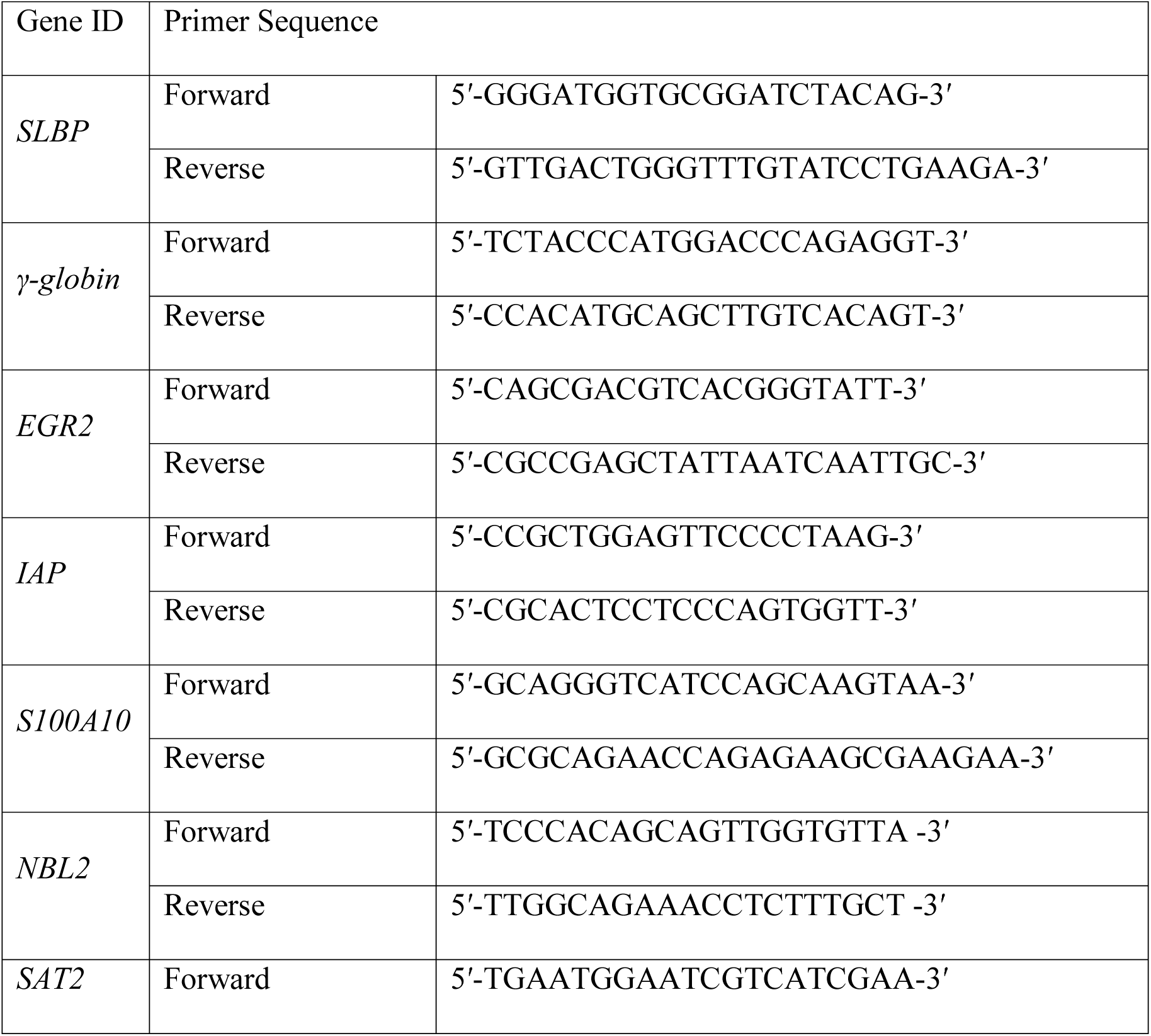
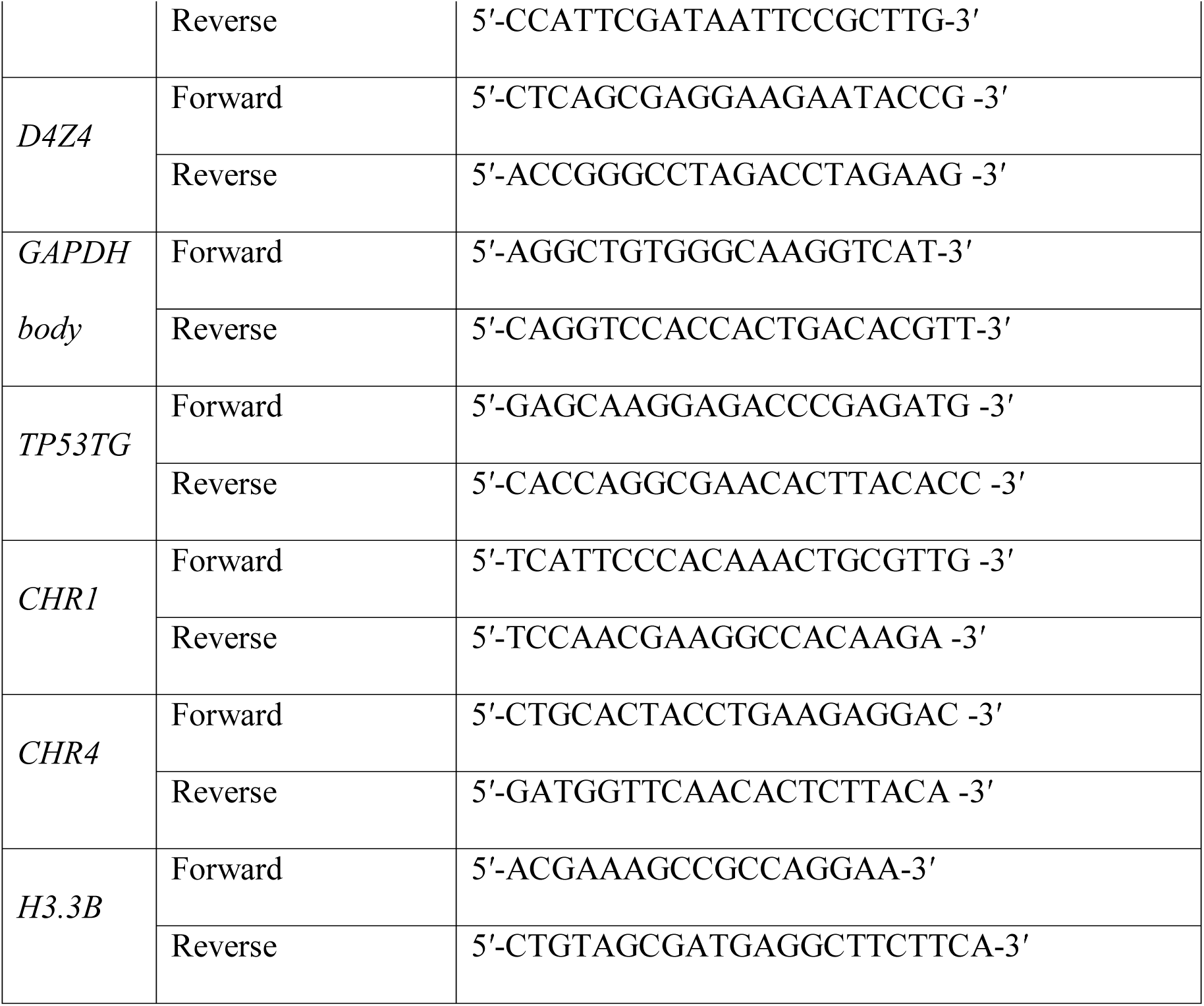

### ChIP-Seq

ChIP-seq libraries were prepared using an Illumina TruSeq ChIP sample preparation kit (IP-202-1024, Illumina, San Diego, CA) according to the manufacturer’s protocol. Sequencing was performed with the IIlumina HiSeq 2500 platform to obtain 50-nucleotide single-or paired end reads. After sequencing, the Illumina reads were mapped to the human genome. Regions of enrichment were identified using MACS peak calling algorithm and GREAT database was used to annotate the aberrant enriched regions.

### RNA sequencing

Total RNA from stable untagged H3.1poly(A) cells was converted into complementary DNA (cDNA) libraries using a Truseq RNA Sample Preparation V2 Kit (Illumina, San Diego, CA). Validation of library preparations was performed on an Agilent Bioanalyzer using the DNA1000 kit. Library concentrations were adjusted to 4 nM, and libraries were pooled for multiplex sequencing. Pooled libraries were denatured and diluted to 15 pM and then clonally clustered onto the sequencing flow cell using the Illumina cBOT Cluster Generation Station and a TruSeq Paired-End Cluster Kit v3-cBot-HS. Sequencing was performed on an Illumina HiSeq2500 Sequencing System using a TruSeq SBS Kit v3-HS. Quality control of FASTQ data was performed using FastQC (version 0.10.1; Babraham Bioinformatics group), and RNA reads were mapped to the human genome (UCSC hg19; February 2009 release; Genome Reference Consortium GRCh37) using STAR (version 2.5.2; Alexander Dobin) (Dobin et al. 2013) with the human reference GTF annotation file (GRCh37). Transcript counts were calculated and normalized using Gfold (version 1.0.8; Jianxing Feng) and DESeq (version 1.6.1; Simon Anders) (Anders and Huber 2010; Feng et al. 2012). The DESeq negative binomial distribution was used to call differentially expressed genes. A total of 2,597 genes were identified as differentially expressed genes (*p* < 0.01). Differentially expressed genes were further investigated for biological function and pathway enrichment using Ingenuity Pathway Analysis (IPA, Qiagen). RNA-seq data have been deposited in the Gene Expression Omnibus database (accession number GSE87541).

### Statistical analysis

Image J image processing software (National Institute of Health) was used to quantify western blotting gel intensities. All statistical significance was calculated and assessed using unpaired, 2 tailed unpaired t-test, where * indicates *p*<0.05 and ** indicates *p*<0.01.

## DATA PROCESSING

### ChIP-seq Data Analysis

Trim Galore (http://www.bioinformatics.babraham.ac.uk/projects/trim_galore/) was used to trim the raw sequence reads ($ trim_galore --phred33 --fastqc --clip_R1 5 --three_prime_clip_R1 3 R1.fq -o OUTDIR). Reads were aligned to the human reference genome (GRCH38/hg38) using BWA [66] ($bwa mem -t 8 INDEX IN.fq > PRENAME.sam). SAM files were then converted into BAM format using samtools [67] ($ samtools view -bS -q 1 -@ 8 PRENAME.sam > PRENAME.bam). Bedtools was used to convert BAM files into BED format ($bedtools bamtobed -i PRENAME.bam > PRENAME.bed). Only 1 copy of the redundant reads that were mapped to the exact same location in the genome was retained. All non-redundant reads from flag-tagged (FH3.3, FH3.3NT3.1, etc.) ChIP-seq experiments were included for downstream analyses without peak calling. MACS2 [68] was used to call peaks under the FDR threshold of 0.01 ($ macs2 callpeak --SPMR -B -q 0.01 --keep-dup 1 -g hs -t PRENAME.bam -n PRENAME --outidr OUTDIR) to identify putative insulators from CTCF ChIP-seq data.

### RNA-seq Data Analysis

RNA-seq data were processed using Salmon[69] ($ salmon quant --gcBias -i INDEX -l A -p 8 -r IN.fq -o OUTDIR). Transcriptome index was built on the human reference genome (GRCH38/hg38). Transcript-level abundance estimates were summarized to the gene-level using the R package tximport[70] for differential expression analysis. DESeq2 [71] was used to identify differentially expressed genes.

### DNase-seq Data Analysis

DNase-seq data were processed using Chilin[72] with default parameters ($ chilin simple -p narrow [--pe] -s hg38 --threads 8 -t IN.fq -i PRENAME -o OUTDIR).

### Composite plot of ChIP-seq profiles at enhancers and insulators

Putative enhancers were identified as top 10,000 shared MACS peaks in two A549 DNase-seq datasets (GSM736580, GSM736506) and are at least 2kb away from any TSS. Insulators were identified as MACS peaks from CTCF ChIP-seq in BEAS-2B cell line (GSM1354438) and are at least 2kb away from any TSS. Each ChIP-seq composite plot covers a region centered at the peak summits (2kb for DNase-seq peaks, 1kb for CTCF ChIP-seq peaks), and average ChIP-seq signals (RPKM) per 20 bps bin were plotted accordingly.

### Data Availability

All sequencing data created within this study are available at NCBI GEO (https://www.ncbi.nlm.nih.gov/geo/) under the accession number GSE135637.

## Supplemental Information

**Supplementary Figure 1. The level of FLAG-tagged H3.3 is not changed by ectopic expression of H3.1poly(A)**

**(A and B)** pcDNA-Empty (EV), pcDNA-H3.1Loop (H3.1Loop), or pcDNA-H3.1Poly(A) (H3.1poly(A)) vector was stably transfected into BEAS-2B FLAG-H3.3 cells separately. (A) RT-qPCR results. Total RNA was extracted from each cell lines. mRNA was converted to cDNA using oligo dT primers. Polyadenylated H3.1 mRNA levels were then measured by quantitative PCR, respectively. Relative mRNA levels were normalized to *Actin* as internal control. * *p*<0.05.

(B) Western blot results. Western blot shows ectopic expression of polyadenylated H3.1 mRNA did not affect ectopic expression of H3.3.

**Supplementary Figure 2. Repeat of ChIP-seq results shown in** Figure 3

**(A-C)** Profile of H3.3-containg nucleosomes across the transcription start sites (TSSs) for 2,000 highly active genes (activate promoters) (A), DNase I hypersensitive sites (enhancers) (B), or CTCF-binding sties (insulators) (C) are shown. In the control cells (blue), H3.3 was enriched at active TSSs, enhancers, and insulators, respectively. The levels were greatly reduced by the ectopic expression of polyadenylated H3.1 mRNA (green).

**Supplementary Figure 3. Cellular functions associated with the top regulator effect network of H3.1poly(A) up-regulated genes identified by IPA**

Illustration of regulator effect networks, such as the aneuploidy, angiogenesis, cell transformation, and cell cycle, associated with up-regulated genes.

**Supplementary Figure 4. RNA-Seq analysis of H3.1poly(A) down-regulated genes**

**(A)** Illustrations of top diseases and disorders associated with 1,437 down-regulated genes.

**(B)** Illustrations of top upstream regulators associated with down-regulated genes.

**(C)** Illustration of top regulator effect network based on down-regulated genes.

**(D)** Illustration of top networks associated with down-regulated genes.

**Supplementary Figure 5. Arsenic induces mRNA polyadenylation of all canonical histones (A-E)** BEAS-2B cells were treated with 1 μM arsenic for 96 hours. Relative polyadenylated mRNA levels of *H2A* (A), *H2B* (B), *H3.1* (C), *H3.2* (D), and *H4* (E) were determined by RT-qPCR with oligo(dT) primers. Values are presented as mean ±S.D. from the experiments performed in triplicate. * *p*<0.05.

**Supplementary Figure 6. Arsenic induces canonical histone protein elevation**

**(A)** Arsenic treated BEAS-2B cells were lysed and immunoblotted against β-actin, H2A, H2B, H3, or H4, separately.

**(B-E)** The bar graphs show relative quantification of canonical histone levels normalized to β-actin. The data shown are the mean ±S.D. from the experiments performed in triplicate. * *p*<0.05.

**Supplementary Figure 7. SLBP overexpression rescues arsenic-induced cell transformation**

**(A)** BEAS-2B cells were stably transfected with pcDNA-empty (EV) or with pcDNA-SLBP (SLBP) plasmid. Western blot was performed using indicated antibodies.

**(B)** mRNA levels of *SLBP* were analyzed by RT-qPCR. Data are mean ±S.D. (n=3).

**(C)** RT-qPCR analysis of polyadenylated H3.1 mRNA level.

**(D)** Soft agar assays. The data shown are the mean ±S.D. (n=3). * *p*<0.05.

