## Supplemental Figures for "Polyadenylation of Histone H3.1 mRNA Promotes Cell Transformation by Displacing H3.3 from Gene Regulatory Elements"

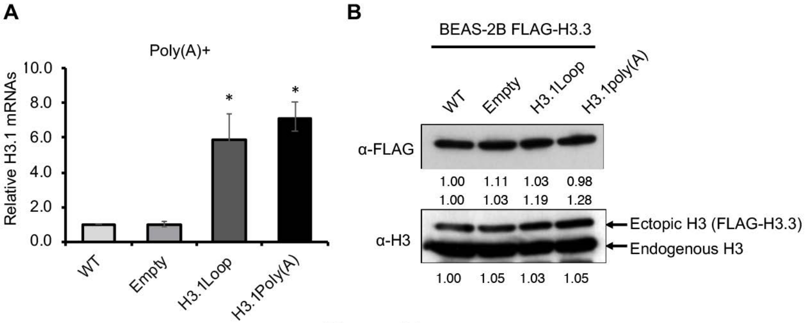
A
Poly(A)+
Relative H3.1 mRNAs
10.0
8.0
6.0
4.0
2.0
0.0
*
*
WT
Empty
H3.1Loop
H3.1Poly(A)
B
BEAS-2B FLAG-H3.3
WT
Empty
H3.1Loop
H3.1poly(A)
α-FLAG
1.00 1.11 1.03 0.98
1.00 1.03 1.19 1.28
α-H3
Ectopic H3 (FLAG-H3.3)
Endogenous H3
1.00 1.05 1.03 1.05

**Figure S1**

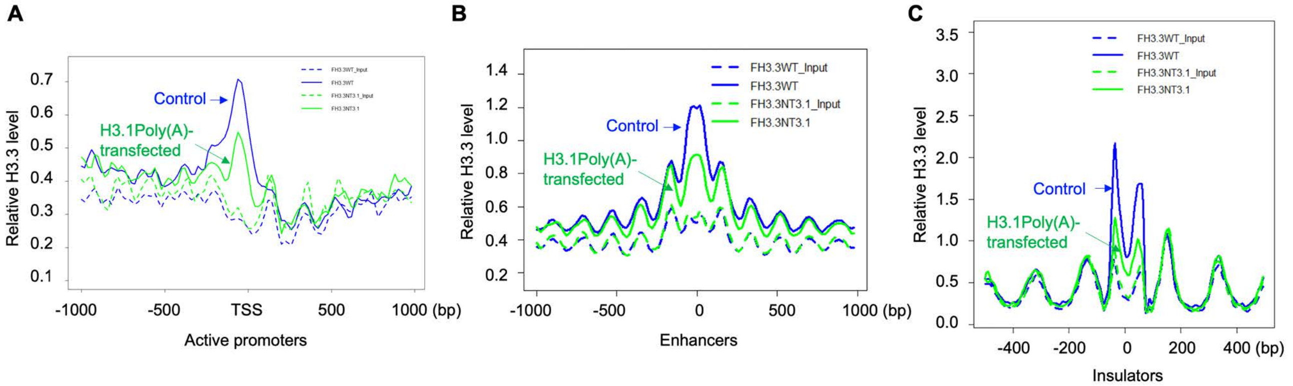

A
B
C
Relative H3.3 level
FH3.3WT_Input
FH3.3WT
FH3.3NT3.1_Input
FH3.3NT3.1
Control
H3.1Poly(A)-
transfected
-1000
-500
TSS
500
1000 (bp)
Active promoters
0
Enhancers
-400
-200
200
400 (bp)
Insulators


**Figure S2**

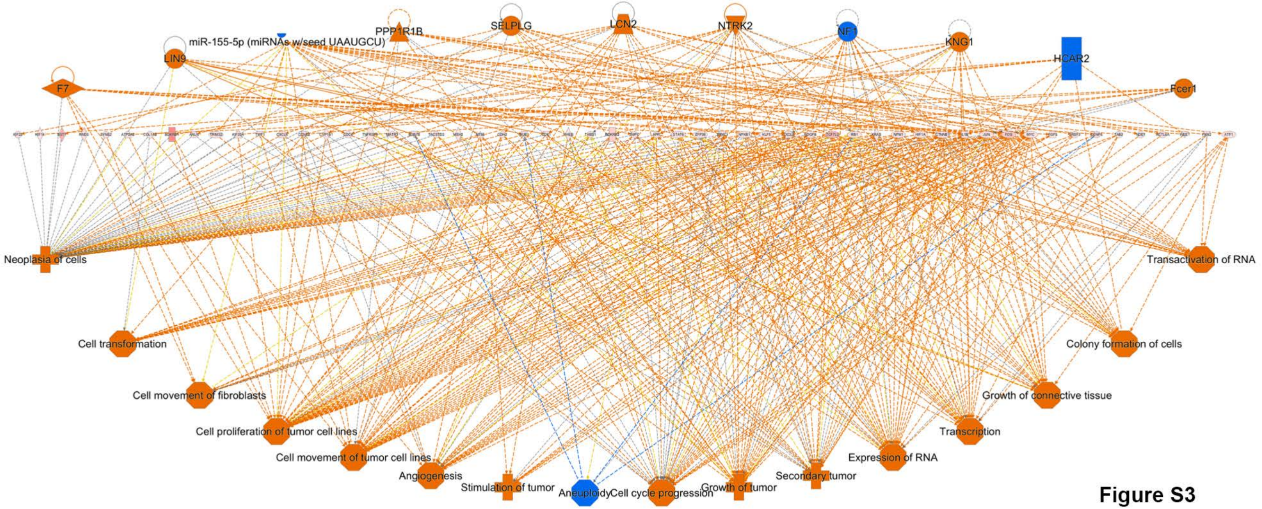
PPP1R1B
SELPLG
LCN2
NTRK2
NF1
KNG1
miR-155-5p (miRNAs w/seed UAAUGCU)
LIN9
HCAR2
F7
Fcer1
Neoplasia of cells
Transactivation of RNA
Cell transformation
Colony formation of cells
Cell movement of fibroblasts
Growth of connective tissue
Cell proliferation of tumor cell lines
Transcription
Cell movement of tumor cell lines
Expression of RNA
Angiogenesis
Secondary tumor
Stimulation of tumor
Aneuploidy
Cell cycle progression
Growth of tumor

**Figure S3**

**A**

| Diseases and Disorders | p-value range | # Molecules |
|---|---|---|
| Cancer | 8.79E-04 - 8.76E-98 | 1330 |
| Endocrine System Disorders | 7.82E-04 - 8.76E-98 | 1212 |
| Organismal Injury and Abnormalities | 1.03E-03 - 8.76E-98 | 1343 |
| Gastrointestinal Disease | 4.49E-04 - 4.45E-48 | 1161 |
| Dermatological Diseases and Conditions | 8.53E-04 - 1.30E-24 | 799 |

**B**

Top Upstream Regulators

| Name | p-value |
|---|---|
| ulipristal acetate | 4.41E-05 |
| TRPS1 | 7.85E-05 |
| ZFHX3 | 1.66E-04 |
| BAIAP2 | 2.59E-04 |
| SH3RF1 | 4.93E-04 |

**C**

Top Regulator Effect Networks

| ID | Regulators | Disease & Functions | Consistency Score |
|---|---|---|---|
| 1 | ETS1,PRDM1,SP1,VEGFA | Abnormality of thymus gland,Arteriosclero... | 8.949 |
| 2 | ISL1,mir-34,NR5A2,POU4F1,PTH1R | Concentration of hormone,Epilepsy (+3 m... | 8.074 |
| 3 | ASCL1,Calmodulin,CTNNB1,DTNBP1,HEY... | Bleeding,Breast or ovarian cancer (+19 m... | 6.629 |
| 4 | CTNNB1,EGR2,IKZF1,WNT3A | Production of hematopoietic progenitor ce... | 6.425 |
| 5 | HEY2,HMGA1 (+6 more) | Bleeding,Breast or ovarian cancer (+9 mo... | 6.155 |

**D**

Top Networks

| ID | Associated Network Functions | Score |
|---|---|---|
| 1 | Cell-To-Cell Signaling and Interaction, Behavior, Nervous System Development and Function | 46 |
| 2 | Cancer, Dermatological Diseases and Conditions, Organismal Injury and Abnormalities | 43 |
| 3 | Developmental Disorder, Skeletal and Muscular Disorders, Connective Tissue Disorders | 41 |
| 4 | Cancer, Organismal Injury and Abnormalities, Hematological Disease | 37 |
| 5 | Cell Death and Survival, Nervous System Development and Function, Small Molecule Biochemistry | 37 |

**Figure S4**

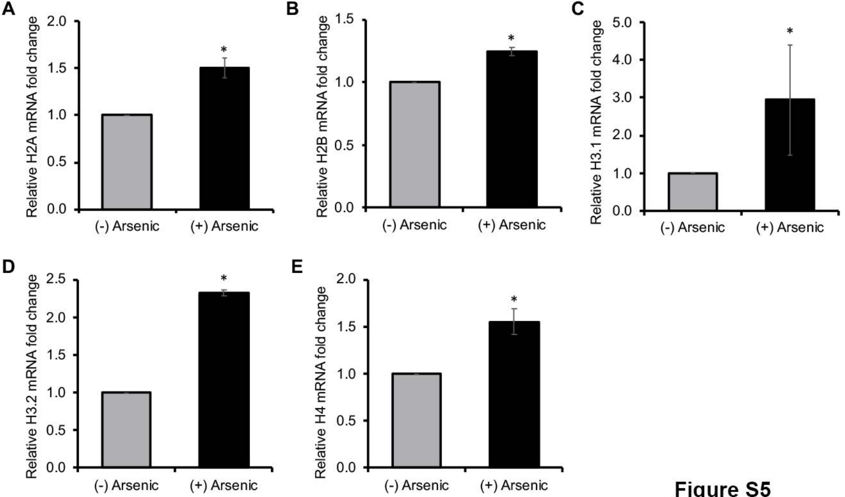

A
Relative H2A mRNA fold change
2.0
1.5
1.0
0.5
0.0
*
(-) Arsenic
(+) Arsenic
B
Relative H2B mRNA fold change
1.5
1.0
0.5
0.0
*
(-) Arsenic
(+) Arsenic
C
Relative H3.1 mRNA fold change
5.0
4.0
3.0
2.0
1.0
0.0
*
(-) Arsenic
(+) Arsenic
D
Relative H3.2 mRNA fold change
2.5
2.0
1.5
1.0
0.5
0.0
*
(-) Arsenic
(+) Arsenic
E
Relative H4 mRNA fold change
2.0
1.5
1.0
0.5
0.0
*
(-) Arsenic
(+) Arsenic


**Figure S5**

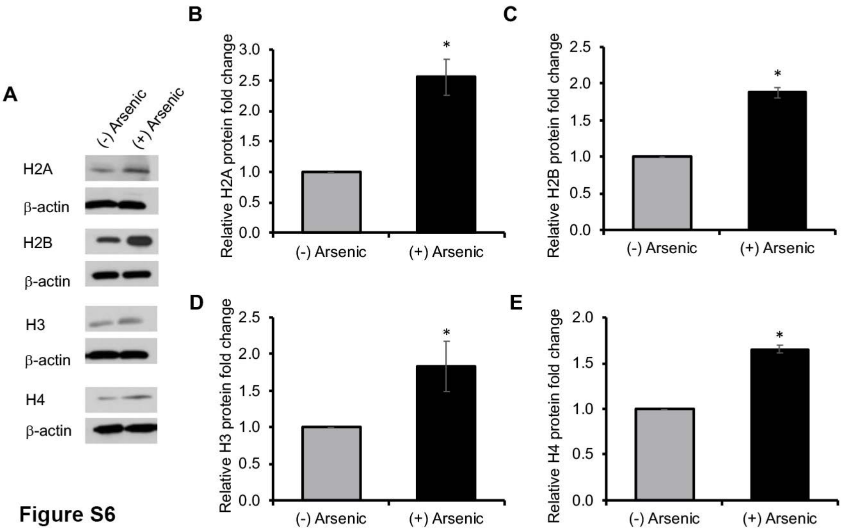
A
(-) Arsenic
(+) Arsenic
H2A
β-actin
H2B
β-actin
H3
β-actin
H4
β-actin
B
Relative H2A protein fold change
3.0
2.5
2.0
1.5
1.0
0.5
0.0
*
(-) Arsenic
(+) Arsenic
C
Relative H2B protein fold change
2.5
2.0
1.5
1.0
0.5
0.0
*
(-) Arsenic
(+) Arsenic
D
Relative H3 protein fold change
2.5
2.0
1.5
1.0
0.5
0.0
*
(-) Arsenic
(+) Arsenic
E
Relative H4 protein fold change
2.0
1.5
1.0
0.5
0.0
*
(-) Arsenic
(+) Arsenic

Figure S6

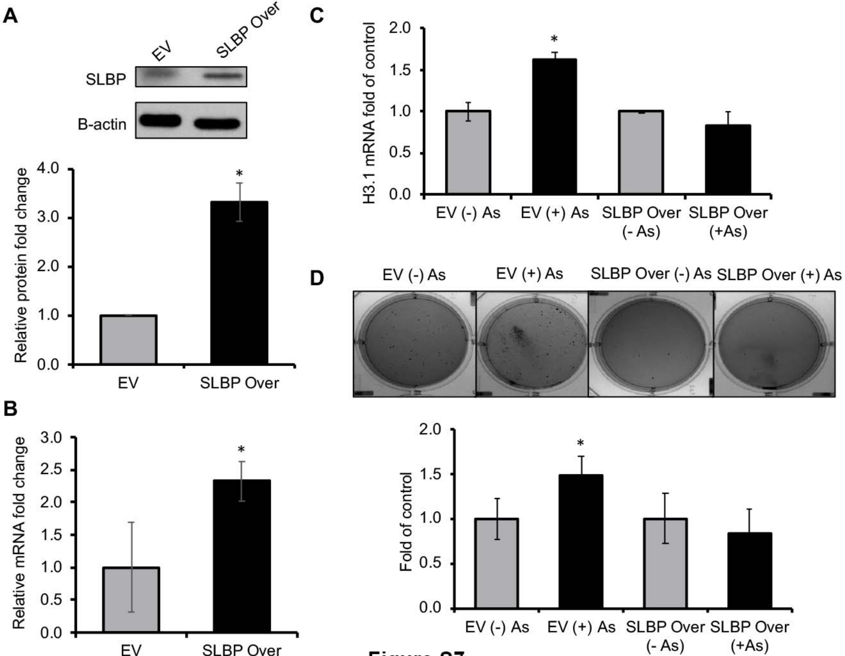
A
EV
SLBP Over
SLBP
B-actin
Relative protein fold change
0.0
1.0
2.0
3.0
4.0
*
EV
SLBP Over
B
Relative mRNA fold change
0.0
0.5
1.0
1.5
2.0
2.5
3.0
*
EV
SLBP Over
C
H3.1 mRNA fold of control
0.0
0.5
1.0
1.5
2.0
*
EV (-) As
EV (+) As
SLBP Over (- As)
SLBP Over (+As)
D
EV (-) As
EV (+) As
SLBP Over (-) As
SLBP Over (+) As
Fold of control
0.0
0.5
1.0
1.5
2.0
*
EV (-) As
EV (+) As
SLBP Over (- As)
SLBP Over (+As)

**Figure S7**
